# Structure-Function in Helical Cardiac Musculature Using Additive Textile Manufacturing

**DOI:** 10.1101/2021.08.18.456852

**Authors:** Huibin Chang, Qihan Liu, John F. Zimmerman, Keel Yong Lee, Qianru Jin, Michael M. Peters, Michael Rosnach, Suji Choi, Sean L. Kim, Herdeline Ann M. Ardoña, Luke A. MacQueen, Christophe O. Chantre, Sarah E. Motta, Elizabeth M. Cordoves, Kevin Kit Parker

**Author notes:** Corresponding Author: Prof. Kevin Kit Parker, 150 Western Ave, Science and Engineering Complex, Boston, MA 02134. These authors are equally contributed (Huibin Chang, Qihan Liu, John F. Zimmerman).

## Abstract

For more than fifty years, it has been hypothesized that the helical alignment of the heart gives rise to its mechanical function. Testing this hypothesis in an engineered environment is difficult, as the fine spatial features and complex three-dimensional (3D) structures of the cardiac musculature are challenging to reproduce using current biofabrication techniques. Addressing this, here we report a new form of additive textile manufacturing, Focused Rotary Jet Spinning (FRJS). FRJS allows for the rapid manufacturing of micro/nanofibers with controlled alignments. Using this method, we manufacture 3D models of the left ventricle, showing that helically aligned scaffolds display increased strain uniformity, axial shortening, cardiac output, and ejection fractions as compared to circumferential models. We then demonstrate how FRJS can enable the assembly of a full-sized model of the human heart’s musculature. This work experimentally confirms that ventricular alignment plays a critical role in ensuring healthy cardiac performance.

## Main Text

The heart is organized in a helical fashion, with cardiomyocytes in the left ventricle smoothly transitioning transmurally from a left-to right-handed helix (*1*). In some cardiomyopathies this organization can be disrupted by maladaptive tissue remodeling (*2, 3*), leading to a more circumferential organization of the myofibrils (*3*–*6*). For more than fifty years, it has been argued that this helical alignment represents a fundamental structure-function relationship which is critical for achieving large ejection fractions (EF) (*1, 7, 8*). *In vivo* studies have provided meaningful insights (*9*–*11*), but testing this assumption in an engineered system requires the precise recreation of these tissue architectures in three-dimensions (3D), with controlled alignments, and micron-scale features.

Approaches using 3D extrusion printing have demonstrated important milestones in biofabrication, including microphysiological devices (*12, 13*), microvasculature systems (*14*) and spontaneously beating heart models (*15, 16*). However, reproducing fine spatial features, while retaining practical production rates, is challenging. This is the result of a fundamental tradeoff. 3D extrusion printing throughput declines rapidly (power law, n=∼2.8) with respect to feature size, while the printing material required scales volumetrically with respect to organ size (**Fig. 1A**). Consider a full-sized human heart. The extracellular matrix components take hours to days to print at current resolutions (∼250 µm feature sizes)(*17*), but at the native feature size of collagen fibrils (∼1 µm), it could take hundreds of years following current trends in scaling (see **Eq. S1**). As these matrix components direct tissue alignment, approaches which can more rapidly produce single micron features are needed, especially while retaining precise control over alignment and organization.

**Fig. 1.**
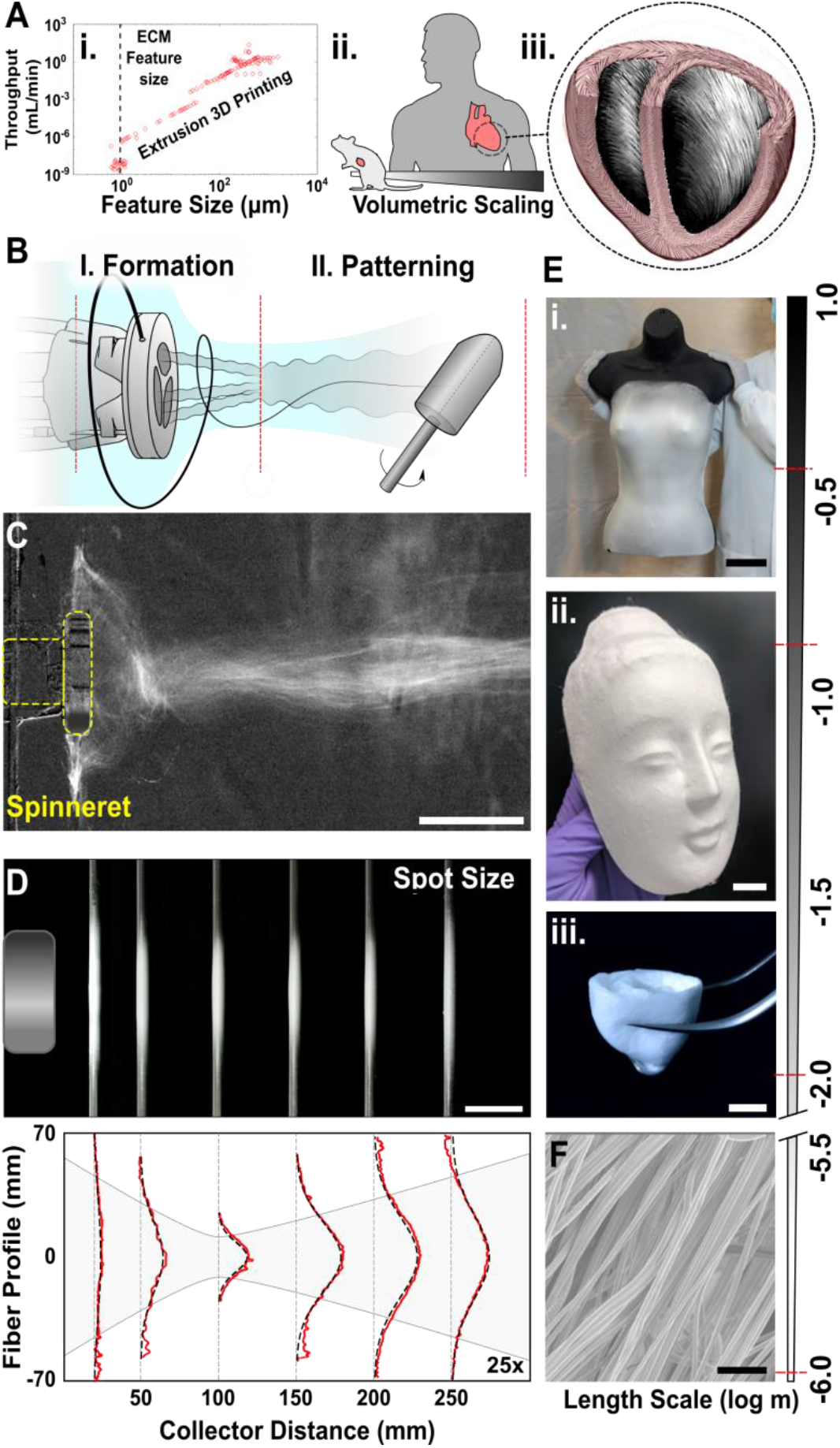
Focused Rotary Jet Spinning. (**A**) 3D printing scales as a power law with feature size (i.), while organs scale volumetrically (**ii**.), while maintaining helical alignments (ventricle schematic, **iii**.), creating a fundamental limit in printing speed. Data collected from literature (*22, 42*–*44*). (**B**) Focused rotary jet spinning separates fiber formation and patterning into two distinct phases, using a focused air stream to structure fibers into confined and aligned stream. (**C**) Differential contrast projection of the fiber stream (maximal projection, scale 5 cm). (**D**) Fiber profiles collected at different points in the stream showing focusing (upper, scale bar, 50 mm), with corresponding thickness distributions (lower) measured as a function of collector distance f(profiles amplified 25x for clarity, red-profile, black-Gaussian fit). (**E**) polycaprolactone (PCL) fiber production across length scales, showing conformal deposition onto a full-scale mannequin (**i**. upper, scale bar, 10 cm), an embossed Buddha mask (**ii**. middle, scale bar, 2 cm), and a rodent-scale ventricle model (**iii**. lower, scale bar, 5 mm). (**F**) Electron micrograph of PCL fibers showing aligned micro/nanofiber formation (mean fiber diameter ∼900 nm, scale bar, 5 µm).

Micro/nanofiber spinning techniques offer a potential solution to this trade-off in reproducing fine spatial features in a high throughput manner. Methods including electrospinning (*18*), melt-blowing (*19*), pull-spinning (*20*), and centrifugal-spinning (*21*), can form micro/nanofibers, some of which offer orders of magnitude greater throughputs in producing single-micron features than 3D extrusion printing (*22*). As a result, these techniques have already been used to engineer tissue constructs, such as heart valves (*23*) and scale ventricle models (*24*). However, unlike 3D printing, fiber spinning approaches often fail to accurately recreate complex 3D geometries with controlled alignments. This lack of precision in fiber organization is due in part to fiber formation and patterning being traditionally interrelated during fiber production. For example, electrospinning requires coupled high voltage electric fields to simultaneously form and pattern fibers (*18*). This process makes it difficult to regulate the orientation of single micron fibers during patterning, while also forming complex 3D structures.

Here we have developed a new form of additive textile manufacturing, focused rotary jet spinning (FRJS), in which we decouple these two processes, first producing fibers using rotary jet spinning and then focusing these fibers into a controlled air stream for deposition. This method allows for the rapid production of micro/nanofibers, while providing control over fiber orientation and alignment. Using this approach, we manufactured both helically and circumferentially aligned models of the left ventricle, showing how these systems are used to probe fundamental biomechanics and scale in such a manner as to conserve the structure-function relationships of the human heart.

### Fiber Manufacture Using Focused Rotary Jet Spinning

In FRJS, decoupling the fiber formation and patterning processes is realized by creating a focused stream of pre-formed fibers (**Fig. 1B&C, Fig. S1, Movie S1**). Rotary jet spinning produces fibers by centrifugal force, pushing polymer solutions through a small orifice in the spinneret. Fibers then undergo jet elongation, resulting in single micron fibers (*21*). This allows for a formation period that is independent of fiber patterning, generating free floating fibers surrounding the spinneret. Next, fibers are pulled into a jet stream blown from the center of the spinneret in a phenomenon known as entrainment (*25*). The entrainment flow is orders of magnitude slower than the air inside the jet (**Fig. S2**), allowing it to minimally perturb fiber formation (**Fig. S3**). Entrainment causes fibers to converge and accelerate towards the jet, which confines and aligns them into a continuous focused stream. To verify this focusing effect, polycaprolactone (PCL) fibers were collected at regular intervals within the airstream to measure deposition profiles. This showed that, at the narrowest point, 95% of fiber deposition occurred within a 5.0 ± 0.3 cm spot size (2σ of Gaussian distribution) (**Fig. 1D**). This resulted in a total throughput rate of 0.1g/min, or 0.03 ghm (grams per hole per minute), which is similar to melt-blowing processes (*19*) but is orders of magnitude greater (∼10^6^) than 3D extrusion printing of single-micron features (**Eq. S1**).

To demonstrate scalability and conformal deposition onto 3D objects, we spun fibers onto targets of differing feature sizes, including a 50 cm female mannequin, a 12 cm Buddha face, and a 1 cm ventricle model (**Fig. 1E, Movie S2**). In each case, targets were coated in less than 30 minutes with micron/nanofibers (**Fig. 1F**), requiring minimal post-patterning modification to achieve conformal deposition (**Fig. S4**). Additionally, we showed that this approach could be used with a variety of material compositions, such as nylon, polyurethane, and gelatin, while still maintaining high throughputs and micro/nanoscale fiber diameters (**Fig. S5**). Collectively, this demonstrated that FRJS is amenable to rapidly manufacturing a large variety of materials and products, spanning multiple length scales in a conformal manner.

### Controlled Fiber Alignment

Next, we examined how FRJS can be used to engineer complex 3D fiber scaffolds with controlled alignments. We hypothesized that the alignment of fibers in the air stream could enable controlled deposition, where tangential collection (θ=0°) should minimally perturb airflow, while head-on deposition (θ=90°) should lead to divergent patterning. To confirm this, the collector angle relative to the fiber stream was modulated during deposition (**Fig. 2A**), with the orientation order parameter (OOP) used as a metric of subsequent fiber organization (*26*). The resulting fibers displayed angle dependent anisotropy, with tangential collection leading to highly anisotropic fibers (θ=0°, OOP = 0.60 ± 0.15), perpendicular collection displaying random fiber orientations (θ=90°, OOP = 0.18 ± 0.03), and intermediate incident angles leading to partially aligned configurations (θ=60°, OOP = 0.50 ± 0.08).

**Fig. 2.**
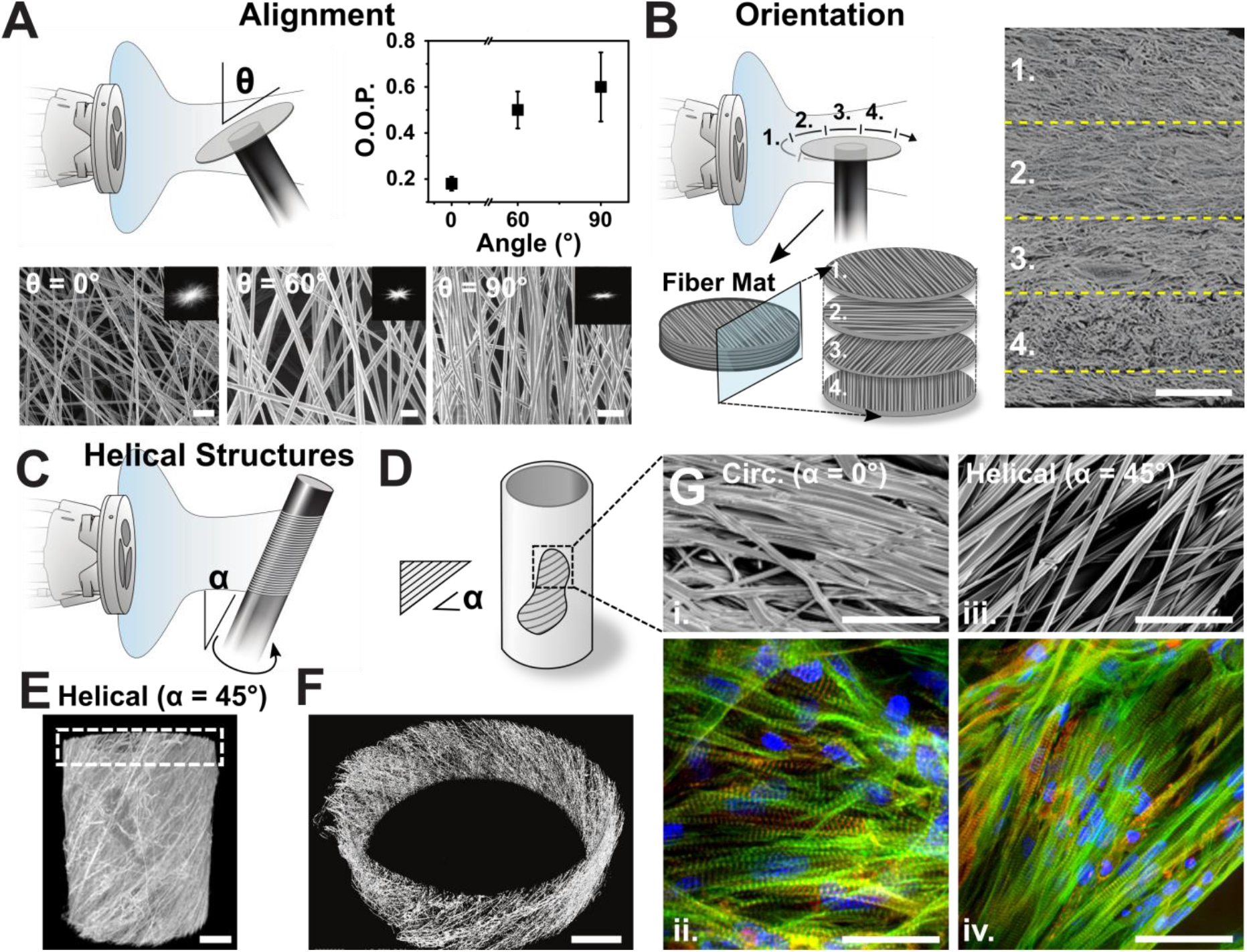
Fiber Alignment Controls Tissue Organization. (**A**) Schematic of fiber collection, showing that the collection angle (θ) dictates the relative alignment of fiber deposition. SEM micrographs (lower)(0°, 60°, and 90°), with corresponding 2D Fourier transforms (inset), indicating the degree of alignment (scale bars, 5μm). Orientation Order Parameter (OOP) indicates the relative alignment across multiple production runs (upper right). (**B**) Rich patterning with controlled orientation can be achieved by manipulating the deposition angle and position of the target (left). Cross section of fiber matt with the corresponding alignments, reconstructed using µCT (right, scale bar, 100 µm). (**C**) Schematic showing fabrication approach for helically aligned cylinders with angle α, and (**D**) corresponding diagram of a released fiber structure, showing the fiber alignment α. (**E**) Reconstructed tomograph of a helically aligned scaffold using µCT (scale bar, 200 µm), with (**F**) inset from highlighted region (scale bar, 200 µm). (**G**) SEM micrograph of gelatin fibers from circumferentially (i., α=0°) and (iii., α=45°) helically aligned cylinders, with corresponding immunofluorescent staining of cardiomyocytes (ii & iv) (NRVM, blue=DAPI, green=f-actin, red=sarcomeres), showing that fibers control subsequent cell infiltration and alignment (scale bars, 50 µm).

As an additive manufacturing technique for nonwoven textiles, this suggested that more complex patterning could be achieved by moving the target relative to the stream. For example, we demonstrate that collecting on an incrementally rotating disk generates a multilayered fiber sheet with controllable fiber orientations as confirmed by X-ray microtomography (µCT) (**Fig. 2B, Movie S3**). Additionally, collecting on an inclined rotating cylinder generates helical alignments (**Fig. 2C-F, Movie S4**). Ensuring FRJS could be used to direct cell infiltration and orientation, gelatin fiber scaffolds were seeded with cardiomyocytes, resulting in highly anisotropic tissues (**Fig. 2G, Fig. S6**). This diversity in manufacturing and ability to direct tissue alignment suggested that FRJS could be used in a hierarchical manner to assist in biofabrication, where the focused air stream provides gross-structural morphology and fibers provide fine structural cues to promote tissue morphogenesis. Having established this technique, we then used FRJS to explore fundamental structure-function relationships in the heart.

### Helically Aligned Model of the Left Ventricle

Cardiac myofibrils are organized helically (*1*), as seen in the cross section of a rodent’s heart (**Fig S7**). Cardiomyopathies such as hypertension (*4, 6*), or mitral valve regurgitation (*5*), can disrupt this myofibril organization leading to the loss of helical alignments and reorganization into more circumferential geometries (*3*–*6*). While the impact of myocardial fiber alignment on cardiac function has been studied *in vivo* (*10, 11, 27*), these studies are characterized by concomitant changes in protein expression and metabolism (*28*). We reasoned that the FRJS’s ability to control fiber alignment could be used to recapitulate these structural changes in the absence of overt biochemical cues, recreating helical models reminiscent of early analytical approaches (*7, 8, 29*). These controlled systems provide a platform for examining how ventricle alignment contributes to cardiac performance and serve as a stepping-stone towards achieving full organ biofabrication.

To test if myocardial fiber alignment led to functional differences in model ventricle performance, we used FRJS to produce both circumferentially (CA, α=0°) and helically (HA, α=30°, 60° with respect to the ventricle’s long axis) aligned gelatin fiber-based model ventricle chambers (**Fig. 3A&B**). HA fiber orientations were selected based on the average myofibril dispersion (34.8°) of a healthy heart (*30*). The scaffolds were then seeded with neonatal rat ventricular myocytes (NRVMs), forming aligned and continuous tissue segments (**Fig. 3C**), which displayed spontaneous contractions after 3-5 days (**Movie S5)**.

**Fig. 3.**
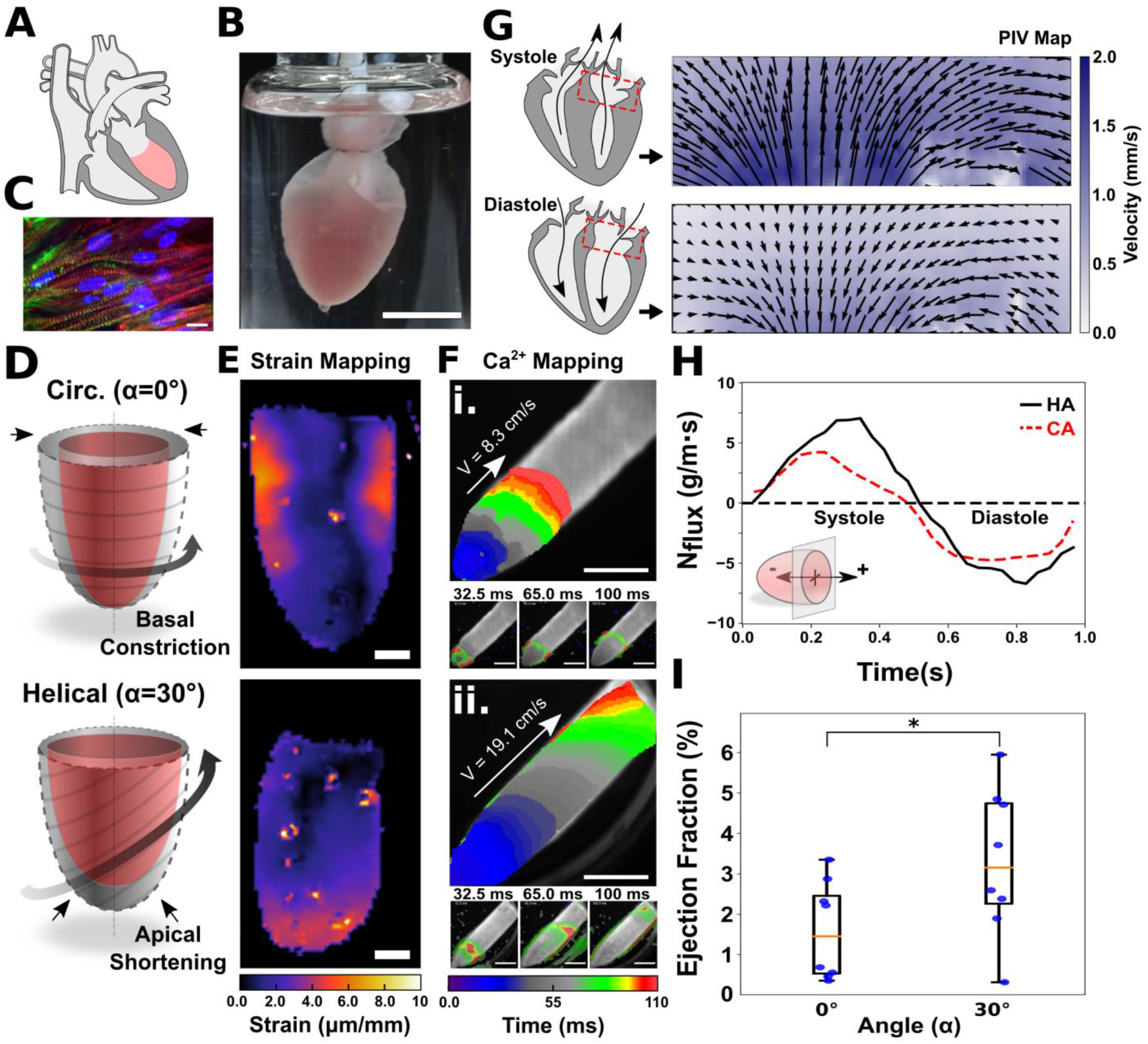
Functional Tissue Engineered Ventricle Models. (**A**) Illustration of the heart. (**B**) Scale model of the left ventricle made with gelatin fiber, seeded with cardiomyocytes (scale bar, 5 mm) (**C**). Immunofluorescent micrograph of cardiomyocytes on the ventricle scaffold (blue=DAPI, green=sarcomeres, red=f-actin, scale bar 10 µm) (**D**) Schematic diagram of a circumferentially (CA, upper) and helically aligned (HA, lower) ventricle, showing differences in wall displacement before (gray outer) and after (red inner) contraction. (**E**) Strain generated during peak contraction in CA (upper) and HA (lower) ventricle models (scale bar, 2 mm). (**F**) Isochrones (upper) with corresponding still-frames (lower), indicating calcium transience along an extended ventricle surface, showing increased transverse wave propagation for HA scaffolds. Point stimulated at apex. (i.-CA, ii.-HA, scale bars, 5 mm). (**G**) Particle imaging velocimetry (PIV) velocity fields taken from the base of a HA ventricle scaffold during peak systole (upper, t=0.3s) and diastole (lower, t=0.7s) (**H**) Representative measurements of the Instantaneous mass flux in the region of interest as function of contraction time. (**I**) Ensemble measurements of ejection fraction (EF) for CA and HA ventricle scaffolds (n = 8 ventricles for each angle, * indicates p < 0.05).

### Ventricle Strain and Deformation

Previously, it has been hypothesized that HA ventricles would display increased apical shortening (*7, 29*), and reduced basal strains (*11, 29*), with a helical organization leading to more distributed strain. To investigate these claims, we examined how model fiber scaffolds deformed during stimulated contraction. Fiber deformation was visualized by submerging cardiomyocyte seeded ventricles into a solution containing fluorescent beads, which non-specifically adhered to the ventricle’s surface. Using digital image cross-correlation, we then performed strain mapping over the ventricle surface (**Fig. 3D, Movie S6**). CA ventricles displayed greater strains near the base of the ventricle, while HA scaffolds showed more uniform deformations, with increased strain at the ventricle’s apex (**Fig. 3E**). Probing this further, changes in the boundary shape were measured using an elliptical fit, observing significant differences in deformation between each alignment. CA samples showed greater basal and minimal longitudinal constriction, while HA ventricles displayed significant apical shortening with reduced basal deformation (n≥5 for each condition) (**Fig. S8**). Overall, these results were consistent with previous analytical and computational models (*7, 29, 31*) indicating more uniform strains, and suggesting that tissue alignment can potentiate changes at the organ scale.

### Alignment Dictates Calcium Wave Propagation

Next, we examined the impact of fiber orientation on calcium wave propagation. In healthy tissues, action potentials propagate faster in the direction of cell alignment, with myocytes showing reduced conduction velocities (CV) in the transverse direction (*32*). Given this biophysical insight, we reasoned that electrical signal propagation may vary depending on the ventricle’s fiber alignment. First, to measure directional signal conduction in tissue scaffolds, we electrically stimulated laminar fiber tissue constructs (point stimulation), and recorded the resulting calcium propagation using an optical mapping system (**Fig. S9**). We observed increased CV in the direction of fiber alignment (longitudinal), with longitudinal (LCV) and transverse (TCV) CVs of 14.9±5.8 and 9.0±3.7 respectively (n = 9 samples) at ratios of ∼1.5-2.0. These values were consistent with previous reports of *in vitro* tissues composed of immature cardiomyocytes where factors such as geometry, calcium handling and gap junction expression can influence CV (*24, 33, 34*). Turning to 3D models, ventricles with extended basal regions – allowing for increased travel distances – were apically stimulated. Isochrones of the calcium propagation revealed that CA scaffolds displayed only modest CVs (8.3 cm/s) along the ventricle’s long axis, while HA ventricles showed greatly increased CVs (19.1 cm/s), with signals taking less than half the time to transverse the ventricle’s surface (**Fig. 3F, Movie S7**). This confirmed that fiber orientation plays an important role in regulating tissue-level calcium propagation, further suggesting the importance of myofibril alignment in excitation-contraction coupling and healthy cardiac function.

### Ventricle Twist

Helical myofibril architectures are also believed to give rise to rotational displacements, or ventricular twist (*1, 35*). In a healthy heart, kinetic energy is stored during contraction in the sarcomeric protein titin and is subsequently released during relaxation (*35*), aiding in diastolic suction. To study if twist was preserved in our model system, we measured rotational displacement at the apex of suspended ventricle scaffolds (**Fig. S10**). Here both HA and CA scaffolds were sutured at the base to a fixed support, allowing the apex to move freely, and were monitored from below during field stimulation. Using edge features to detect rotation, we observed that CA scaffolds showed minimal torsion (1.35 ± 1.1°, n=7). Conversely, HA scaffolds showed an approximately four times greater rotational displacement (5.4 ± 3.6°, n=7) (**Movie S8)**, representing to our knowledge the first report of ventricle twist in an *in vitro* system. Combined, these changes in deformation, calcium propagation and rotational displacement, indicated substantive structural differences across scaffold orientations, suggesting the potential for functional differences based on fiber alignment.

### Cardiac Output and Ejection Fractions

To determine if these structural changes in alignment and deformation lead to altered function, we then used cardiac output and ejection fraction (EF) as quantitative metrics of cardiac performance. To measure these values we first monitored pressure-volume (PV) changes in the ventricle scaffolds using catheterization, observing the formation of complete oblate PV loops (**Fig. S11**). However, submerged gelatin fibers are a poor dielectric, making it difficult to obtain consistent results in our synthetic scaffolds using conductance catheterization. Consequently, particle imaging velocimetry (PIV) was used to further quantify cardiac performance (*36*).

Here, ventricle constructs were suspended in a bath containing neutrally buoyant fluorescent beads, and bead displacement was tracked. This allowed for the construction of two-dimensional (2D) velocity fields surrounding the basal opening, as shown for a HA ventricle construct (**Fig. 3G**). Using PIV, we then evaluated the instantaneous mass flux resulting from ventricle contraction (**Fig. S12, Movie S9**), observing cyclic outputs, with fluid being expelled during systolic contraction, and refill occurring during diastole (**Fig. 3H**). Summing the fluid displacement over systole yields the total cardiac output, with values of 1.3±0.9 g/m and 2.6±1.3 g/m (n=8 each) for CA and HA ventricle scaffolds, respectively (**Fig. S12G**). This represented a significant (p < 0.05) increase in cardiac output based purely on ventricular tissue alignment. Next, normalizing the cardiac output by diastolic ventricle volume and fluid density we obtained estimates for the resulting EFs. This yielded average EFs of 1.6 ± 1.1% and 3.3 ± 1.7% in the CA and HA case, respectively (n=8 each), with a maximum EF of 5.9% observed in the helical case (**Fig. 3I**). Although significantly lower than *in vivo* EFs (∼60-80%), these values are consistent with previously reported *in vitro* systems (*24*)(*37*), and are expected given the limited tissue depth. With a ∼100% increase in EFs for helical architectures, these results indicate that tissue alignment plays a critical role in modulating ventricular ejection fractions.

### Dual Chamber Ventricles and Full-Scale Heart Models

Moving towards higher fidelity representations of the human heart, we then used FRJS to create models with multiple helical angles, including a dual-chambered ventricle (DCV) and a full-scale human heart musculature model (**Fig. 4A**). DCVs were manufactured using a multistage process, creating inner and outer helical layers, with an intermediate circumferential sheet reminiscent of native myocardium (**Fig. S13, Fig. 4B, Movie S10**). These helical structures were confirmed using µCT, indicating three distinct transmural layers along the septal wall (**Fig. 4C&D, Movie S11-12**). Showing that DCVs could support human-derived cardiac tissues, scaffolds were then seeded with human induced pluripotent stem-cell derived cardiomyocytes (hiPSC-CMs), which were differentiated *in vitro*, forming visible spontaneous contractions after seven days, and staining positive for sarcomeric α-actinin (**Fig. S14**). Examining an excised segment of the left-ventricle we observed that cells adhered to the fibers, forming aligned tissues (**Fig. 4E**), wherein infiltration was primarily restricted to the scaffold surface owing to an absence of vascularization. Additionally, calcium imaging revealed sustained wave propagation across the ventricle surface, occurring primarily in the direction of fiber alignment, indicating the formation of a continuous cardiac syncytium (n=2) (**Fig. 4F**). Overall, this indicated that fiber constructs can also support human stem cell derived tissues in complex geometric geometries and controlled alignments.

**Fig. 4.**
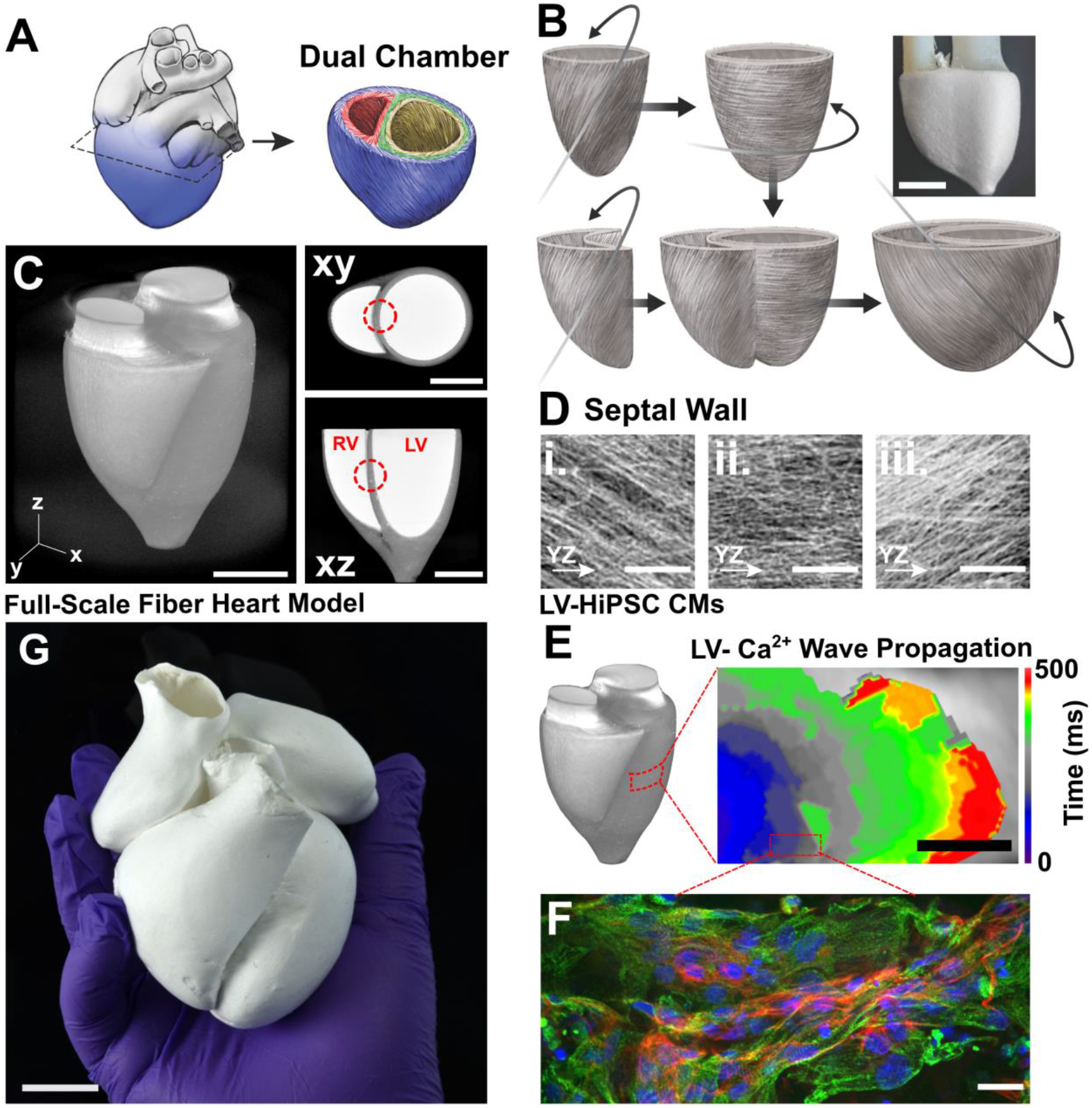
Multiscale Heart Models. (**A**) Simplified design of a tri-layered, dual chamber ventricle (DCV) mimicking the native ECM alignment of the heart. (**B**) Four-step manufacturing of DCV model, with image of the resulting structure (scale bar, 5 mm). (**C**) µCT imaging of the DCV design, with corresponding cross-sections (scale bars, 5 mm). (**D**) High magnification µCT images taken from the intraventricular septum (highlighted in red), showing tri-layer alignments from the respective regions (insets) (scale bars, 25 µm). (**E**) Isochronal map of calcium wave propagation in human induced pluripotent stem cell-derived cardiomyocytes (hiPSC-CM) across an excised portion of the level ventricle, showing anisotropic propagation (scale bar, 2 mm). (**F**) Immunofluorescent micrographs of taken from an excised region of the DCV’s left ventricle wall, showing aligned cardiac tissue (blue=DAPI, green=sarcomeres, red=f-actin, scale bar, 100 µm). (**G**) Full-scale four chambered human heart model composed of single micron/nanofibers (scale bar, 2 cm).

To demonstrate the scalability of FRJS with regards to biofabrication, a full-sized acellular model of the human heart musculature was constructed. This was assembled by patterning micro/nanofibers onto dissolvable collectors in the shape of each of the heart’s four chambers (**Fig. S15**). Individual chambers were then connected by chemical annealing, before dissolving the interior supports, resulting in a standalone scaffold construct (**Fig. 4G**). While full-sized anatomical models have previously been produced using thermoplastics and hydrogels (*17*), these systems typically lack the micron-scale features needed to direct myocyte alignment. Here, using a micro/nanofiber-based approach, these local structures can be preserved across entire tissue volumes, allowing for the hierarchical assembly of tissue constructs. While there is still work needed to generate tissues containing billions of cells with diverse populations (e.g. neurons, vasculature, fibroblasts, etc), as found in full-sized organs, our proof of concept 3D organ models demonstrate that FRJS scaffolds support human-derived tissues and allow for rapid assembly of full-sized models of the musculature. These key features mark fiber-based manufacturing as a promising approach for achieving whole organ biofabrication, which can be used as an alternative to or in conjunction with emerging biomanufacturing platforms such as 3D printing.

## Discussion

Here we have shown a novel approach for biofabrication using FRJS to overcome trade-offs in spatial resolution and throughput for manufacturing complex 3D structures. This allowed for the rapid assembly of functional ventricle models capable of recapitulating emergent phenomena *in vitro*, including ventricular twist, strain displacement, and myocardial angle-dependent ejection fractions. While further work is needed to achieve full-scale *de novo* organ fabrication, our model systems provide clinically relevant metrics and offer the potential for improving the predicative capabilities of biomedical assays for organs where tissue alignment is used to regulate fluid displacement. Demonstrating this, we have shown how cardiac myofibril alignment regulates ventricular performance, confirming theoretical predictions made over fifty years prior (*7*). These results suggest that a reduction in helical architecture can alter cardiac performance, reminiscent of some pathological cases such as the increased circumferential strain (*38*) and reduced ejection fractions observed in hypertensive patients (*4, 6*). This lends support to emerging clinical approaches using conical ventricle reshaping to address heart failure (*39, 40*). Although ventricle reshaping has been viewed as controversial, with a landmark surgical study showing no significant increase in survival or functional outputs (*40*), subsequent reports have argued this was the result of surgical reshaping that focused on volume rather than shape (*39, 41*). The present study lends credence to this idea, and further suggests that special consideration should be given to the final tissue alignment in post-reconstructive surgery.

In addition to biofabrication, FRJS may also serve an important role in other additive manufacturing applications, where a material’s properties are determined by its microstructure and alignment. The high surface area to volume ratio of nanofibers makes them ideal candidates for a variety of industrial applications, such as air filtration systems, face masks and drug delivery. Additionally, FRJS lends itself to applications requiring well controlled fiber alignment and anisotropy, which have been difficult to achieve using traditional fiber spinning approaches. In this regard, FRJS provides a high-throughput option for obtaining high quality fibers with a variety of material compositions, controlled fiber alignments, and complex geometries.

## Supporting information

Movie S1 - Fiber Stream

Movie S2 - Fiber Deposition

Movie S3 - Multilayer Alignment

Movie S4 - Helical Structures

Movie S5 - Ventricle Contraction

Movie S6 - Ventricle Strain Mapping

Movie S7 - Calcium Propagation

Movie S8 - Ventricle Twist

Movie S9 - Cardiac Output

Movie S10 - DCV Manufacture

Movie S11 - DCV Scan

Movie S12 - DCV High Mag Septal wall

## Author Contributions

K.K.P. supervised the research. K.K.P., H.C., Q.L., J.F.Z, and L.A.M., designed the study. Q.L. and H.C. designed and produced the fiber spinning platform. H.C and J.F.Z, manufactured, cultured and performed the ventricle experiments. K.Y.L performed optical mapping experiments. H.C., Q.L., J.F.Z., and K.Y.L analyzed the data. Q.L. performed simulations. L.A.M, C.O.C, S.E.M, G.T, and E.M.C. performed additional supporting experiments. S.L.K., H.A.M.A. S.C. and Q.J. helped with animal protocols and stem cell culture to obtain cardiomyocytes. M.M.P. and H.C. performed SEM experiments and analysis. M.R., H.C., J.F.Z, and C.O.C produced the full-sized heart model. All authors discussed the results and contributed to the writing of the final manuscript.

## Disclosures

Harvard University filed for intellectual property relevant to this manuscript, listing J.F.Z, Q.L. H.C., and K.K.P. as inventors.

## Acknowledgments

The authors would like to thank Dr. A.G. Kleber, M.D. for discussions regarding cardiac physiology, P. Campbell for rodent heart isolation, W.T. Pu, M.D. for providing hiPSC cell lines, H-Y.G. Lin for his assistance with micro-CT and the Qingzhou Museum (Qingzhou, China) for sharing 3D models of the Buddha head (Longxing Temple, 5th century A.D). H.A.M.A. would like to thank the American Chemical Society for support through the Irving S. Sigal Postdoctoral Fellowship. This work was sponsored by the John A. Paulson School of Engineering and Applied Sciences at Harvard University, the Wyss Institute for Biologically Inspired Engineering at Harvard University, the Harvard Materials Research Science and Engineering Center (DMR-1420570, DMR-2011754), the National Institutes of Health with the Center for Nanoscale Systems (S10OD023519) and National Center for Advancing Translational Sciences (UH3TR000522, 1-UG3-HL-141798-01). The content is solely the responsibility of the authors and does not necessarily represent the official views of the National Institutes of Health.

