## Supplemental Methods and Material for "Structure-Function in Helical Cardiac Musculature Using Additive Textile Manufacturing"

**This file includes:**

Materials and Methods
Figs. S1 to S15
Equations S1 to S4
Captions for Movies S1 to S12

**Other Supplementary Materials for this manuscript include the following:**

Movies S1 to S12

### Materials and Methods

3D Extrusion Printing Speed Estimation

To compare manufacturing approaches for biofabrication, we estimated the time required to assemble a full-scale human heart model using dimensional analysis and current trends in 3D extrusion printing such that

$$t = W_o \cdot P_o \cdot ECM_{\%} \cdot V_e(s) \quad \textbf{(S1)}$$

where t is the total print time, $W_o$ is the weight of the organ, $P_o$ is the organ's density, $ECM_{\%}$ is the weight percent composition of extracellular matrix proteins for a given organ, and $V_e(s)$ is the printing method's extrusion speed based on the specific feature size, s. This model assumes perfectly optimized printing paths, no retracing of steps, and continuous material extrusion. For an adult heart, we assumed an organ weight of 280 g, a material density of 1.06 g/mL, and a collagen ECM composition of 1.2% (*45*). With regard to $V_e(s)$, to our knowledge no examples of single micron 3D extrusion printing have been reported. However extrusion rates can be estimated using a linear regression of current trends in 3D extrusion printing (Fig 1.A), yielding an estimate of ~$4.57 \times 10^{-8}$ mL/min for single micron features (~$4.84 \times 10^{-8}$ g/min)(*22*, *42–44*). For a full-scale heart model, this suggests it would take a total print time of ~1104 years to reproduce every feature with single micron fidelity, or ~137 years to reproduce only the collagen ECM components. To calibrate this model, we also estimated total print times for known literature values. For instance, printing with 250 µm feature resolution, we estimated it would take ~14.5 hours to print a full weight human heart model. This estimate was in line with (order-of-magnitude) examples presented by Mirdamadi et. al., who reported a total print time of ~92 hours (*17*).

Focused Rotary Jet Spinning Construction and Setup

The basic design of the focused rotary jet spinning system is shown in Fig. S1A.

The system consisted of a high-speed motor spindle (NR-4040, part No. NSK 9213) purchased from Nakanishi, with the following related parts: collection chuck (CHK-6.35AA, part No. NSK 91601), collect nut (part No. NSK 2158), control unit (E3000, NE 211, part No. NSK 9775), motor cord (EMCD-3000J-4M, part No. NSK 1768), air-line kit ( AL-A1205, part No. NSK 4505), and controller extension (E3000-PEX4, part No. NSK 8407). The air blower that was used to form the focused fiber stream was 3D printed from an Objet30 3D printer (Stratasys, Eden Prairie, Minnesota, USA) using VeroWhitePlus® (RGD835) photopolymer resin, Fig.S2B. The air blower was press-fitted onto a motor spindle and is connected to the building's compressed air source through an air pressure regulator (part No. NSK 4505, Nakanishi). The spinneret is machined from aluminum 7075, and is 6.4 cm in diameter, with the spinneret design containing three orifices, or holes, each 400 µm in diameter. The spinneret lid is machined from polypropylene for its solvent resistance, and is press-fitted onto the bottom to ensure proper sealing.

Polymer solutions were fed into the spinneret through a small needle, which was fit in place into a small hole in the rear of the air blower. Using an automated syringe pump, purchased from Harvard Apparatus (part No. 703007), polymer solutions were passed from a syringe (60 mL, Luer-lok™ Tip) to the needle (BD company) via flexible tubing, feeding polymer solution to the spinneret. The Polyfluoroalkoxy alkane tubing was purchased from Saint-Gobain (Part No.

TSPF35-0125-031-50). The luer lock couplings (part No. 51525K326 and part No. 51525K24) was obtained from McMaster-Carr company. Universal stand (part No. 570-12000-00) and clamps (part No. 36300550) holding the motor were purchased from Heidolph Instruments. For fiber collection, the collection mold was fixed into a high-speed motor spindle (NR-4040, part No. NSK 9213) with a Nakanishi E4000 controller.

Solution preparation and fiber spinning

Polycaprolactone, PCL: Polycaprolactone (PCL, average Mn=80,000, Sigma-Aldrich 440744) was used as the default polymer in this study, unless otherwise specified, while hexfluoroispropanol (HFIP, Oakwood Chemical 003409) was used as the primary solvent, unless otherwise specified. Polymer solutions were dissolved in HFIP at room temperature on a magnet stir plate (PC-4200 Corning) overnight. Concentrations of 6g/100mL PCL solution, 6g/100mL Nylon 6 pellets (Sigma-Aldrich 181110), 8g/100mL polyurethane (Lubrizol, Pellethane 2363-80A) and 5g/100mL Gelatin type A (Sigma-Aldrich G2500) were used in the corresponding experiments.

Polymer fiber preparation method**:** For fiber spinning, rotation speed of the spinneret is set to 10,000 RPM, with a pressure input of 0.2 MPA to the air blower, unless otherwise specified. The number of orifices, or holes, in the spinneret, solution flow rates, total solution volumes and collection distance are described in each experiment. All collection targets are treated with Teflon spray (DuPont, Teflon Non-Stick Dry-Film Lubricant) before deposition. Two of the three orifice on the spinneret were used in spinning unless otherwise specified. Here, a Kapton tape (1/4" Wide, 15' Long, 0.0025" Overall Thickness, part No. 7648A731, McMaster-Carr) was used to block the orifice. Polymer solutions were fed into the spinneret at 0.4mL/min unless otherwise specified.

Deposition on female mannequin, Fig.1D: Female molded black shirt form (part No. 70211) was purchased from Amazon. Fibers were directly deposited on female shirt mannequin. 200 mL PCL solution at 6g/mL concentrations were spun at 2 mL/min feed rates, using all 3 orifices on the spinneret. The target was placed about ~30 cm from the spinneret, while the spinneret was held by hand and manually manipulated to uniformly spray fibers onto the target.

Deposition on Buddha mask, Fig.1E-G: 3D model of the Buddha's face from a 5$^{th}$ century statue is obtained from Qingzhou Museum in China. The 3D model is printed using an Objet30 printer (Stratasys, Eden Prairie, Minnesota, USA) with the VeroWhitePlus® (RGD835) photopolymer resin. A negative mold of the Buddha face is made by Mold Max™ 60 (Smooth-On), where parts A & B are mixed 100A & 3B by weight and cured for 24 hours. The negative mold was then later used for embossing. PCL fibers were directly deposited on buddha face mold. Spinning conditions were the same as those used for the female mannequin, using a total of 20 mL of solution. To deposit fibers, the Buddha face was held by a clamp and the spinneret was manually manipulated to uniformly deposit fibers. After spinning, the negative mold is immediately pressed onto the fiber deposition, to emboss the face's features.

Fiber sheet with rotating alignment, Fig.2D: A 6 cm in diameter flat circular collector was cut from an acrylic sheet (1/16" thickness, part No. 8560K171, McMaster-Carr) using an Epilog Mini 24 laser cutter. The collector was taped onto side of a 10 mm steel bar, which was inserted into the fiber stream, minimally perturbing air flow. The steel bar was inserted into the chuck of the spindle motor in the Y axis, and the chuck was rotated every 5 minutes to obtain scaffolds with multiple angles. In total,16 mL of polymer solution was collected at a distance of ~8 cm.

Fiber tube with helical alignment, Fig.2E: A low contrast stiff Nylon MicroSwab (TX730, ITW Texwipe) with 0.9 mm diameter was used as the collector. The collector was maintained at a helical angle of 45°, and was rotated at 2,000 RPM, with the following spinning conditions: 0.4 mL/min flow rate, ~ 1.2 mL PCL solution, and ~20 cm collector distance.

Single ventricle scaffolds: A 3D printed target of the left ventricle measuring ~10 mm in diameter was used. ~5mL gelatin solution of 5g/mL concentration was deposited onto the target at a ~20cm collection distance, with rotation speeds of 2,000RPM. The target is air dried for 1 hour to remove residual solvent. The gelatin scaffold was then immersed in 100mL ethanol/water (95/5 vol%, 200 proof ethanol VWR and DI water) solution containing 0.96 gram of 1-ethyl-3-(3-dimethylaminopropyl)carbodiimide (EDC, Sigma-Aldrich), and 0.23 gram of N-hydroxysuccinimide (NHS, Sigma-Aldrich) for 24 h for cross-linking. Then scaffold was then washed by PBS three times to remove unreacted chemicals, and then stored in de-ionized water. Prior to culture, the scaffold is then sterilized in 70% ethanol for 30 minutes and placed under UV Ozone for at least 7 minutes. Scaffolds were then soaked in 50 µg/mL human fibronectin (BD Biosciences) dissolved in PBS for 30 minutes at 37° C prior to seeding.

Fibrous dual-ventricle model, Fig. S13: Targets with the shape of left and right ventricle are designed in SolidWorks and are 3D printed. The multi-layered alignment was achieved through a 4 step fabrication process, as shown in **Fig. S13**. First, A 3D printed target in the shape of the left ventricle was positioned at a 45° angle relative to the fiber stream. The target was then rotated counterclockwise at 2,000RPM to collect a helically aligned inner layer. The target was then reoriented vertically, 0° relative to the fiber stream, and spun at 10,000RPM to collect the circumferentially aligned middle layer. Next, a separate 3D printed target with the shape of the right ventricle was positioned at a 45° angle relative to the fiber stream. The target was then rotated counterclockwise at 2,000RPM to collect a helically aligned inner layer for the right ventricle. The left and right ventricular targets were then pressed together with and positioned within the fiber stream at an angle of 45º. The combined targets were then rotated clockwise at 2,000RPM to collect the helically aligned outer layer. For each layer, ~2.5 mL PCL or gelatin solution was deposited and collection distances of ~20 cm were used. For the steps to make helical structures, an additional collector was used to prevent the accumulation of fibers on the edge of the scaffold.

Full-sized Four-Chamber Human Heart Model

Reconstruction of Digital Heart Chamber Topology and 3D Printing, Fig. S15A-C: An opensource scan of a life-sized human heart was acquired from BodyParts3D (https://lifesciencedb.jp/bp3d/), and the heart's musculature was isolated from the non-muscular components. To enable clean 3D printing, in the absence of polygonal defects, a topologically matching model was then built for each chamber, conforming to the surface of the original scan. Cavities between chambers resulting from scanning imperfections were topologically filled, to facilitate full connection between heart chambers. Reconstructed heart chambers were then printed at scale using polylactic acid (PLA) as a base polymer.

Silicone Chamber Molds, Fig S15D: To create mold containers, a laser cutter (Epilog Mini 24) was used to form acrylic rectangular walls of sizes 5.25”x5.25”, 4.125”x4.125”, 3.125”x3.125”, and 4.125”x3.125”, with a depth of 0.25”. Differently sized containers were then created to match the varying sizes of each of the four PLA heart chamber models, with each container consisting of four walls without a top or bottom. The bottom of each container was then constructed from a layer of plasticine (oil-based non-hardening clay). The printed model chambers were then placed on top of the plasticine floor, with the cavity situated as to minimize air pocket formation during molding. The midsection of each chamber was then marked off with graphite, and a plasticine seal was constructed at the midsection level to seal off half of the acrylic container. The gaps between acrylic walls were sealed off using tape and parafilm.

For forming the molds, a silicone rubber base and activator (Cast-a-Mold) were thoroughly mixed at 1:1 ratios by volume. During the subsequent 40 minutes of working time, mixtures were degassed for 15 minutes in a vacuum chamber. The silicone mixture was then poured into the mold container, filling the half of the container not blocked by plasticine. The silicone was then left to harden overnight, and the silicone was then separated from the acrylic container, and the plasticine cleaned off. The half silicone mold was then sprayed with Teflon spray to prevent adhesion, and the molding process was then repeated for the second-half of the chamber. The next day the silicone mold was removed from the container, and the two halves of the silicone mold were then separated from each other using an initial incision. The original reconstructed PLA heart chamber was then removed from the mold, and the mold was tested for its ability to contain liquid without leaking. As a final step, three holes were drilled in each silicone mold: a 0.5” hole for pouring the sugar solution; a 0.5” hole for inserting a steel rod used to hold and rotate the heart chamber; a .125” hole near the pouring hole to facilitate air escape.

Dissolvable Sugar Molds, Fig S15E: Positive dissolvable sugar molds were then formed for each chamber, using the negative silicone rubber molds. Here, powdered cane sugar was first dissolved in excess in water, and then brought to a boil at temperatures of 145° C, removing the water and resulting in molten sugar. While still hot, sugar was then poured into each of the heart chambers and allowed to sit for at least an hour before being removed from the molds. Prior to pouring, steel rods, ~10 mm in diameter, were inserted into each chamber to act as supports for spinning. Where possible, rods were inserted at the location of arteries or valves, to preserve the overall morphology.

Micro/Nanofiber Formation, Fig S15F-J: For each component, a 120 mL solution of polycaprolactone (PCL) (6g/mL concentrations) was spun onto the sugar molds at a 2 mL/min feed rate, using all 3 orifices on the spinneret. The targets were placed about ~20 cm from the spinneret. Each chamber was positioned at an angle of 45° relative to the fiber stream. The targets were rotated counterclockwise at 2,000RPM to collect helically aligned layers along the chamber surface.

Chambers were then connected by chemical annealing of the exterior fiber shell, spraying the surface with a light mist of HFIP. Chambers were then placed together for ~30 seconds before they were bonded together. The heart was then submerged in de-ionized water overnight, dissolving the interior supports, resulting in a standalone scaffold construct.

Fiber distribution inside the stream

The focus of the fiber stream is characterized by collecting fiber on a thin steel rod at different distances (**Fig.S5**). The steel rods are 10 mm in diameter and rotate at 2000RPM. ~8mL PCL solution is deposited on each sample at 2mL/min feed rate using 3 orifices. Samples were then imaged against a black background using a Nikon camera (Nikon D750) at a pixel resolution of 6016x4016. The diameter of the steel rod diameter is used to calibrate the size of pixels (~0.031 mm/pixel). During the image processing, we draw a straight baseline between the naked top and bottom parts of the rod. The straight line marked the position of the rod underneath the fiber deposition. The image is then turned to black and white image use Huang threshold filter. The distance between the boundary of the fiber layer and the baseline is then collected as the thickness of the fiber layer.

For each sample, the thickness profiles were taken from 4 different angles. 3 samples were collected for each distance. Each thickness profile was fitted with a Gaussian profile use the curve fitting tool in Matlab. For each collection condition, the standard deviation $\sigma$ is averaged. The averaged $\sigma$ is fitted with linear relation with respect to distance with a constrained slope of $d\sigma/dX = 0$.

Differential contrast imaging, used to increase the contrast of the fiber stream (**Fig. 1C**, **Video S1**), was obtained by subtracting temporally adjacent video frames from one another. This highlights the difference in frames, rather than the absolute value increasing contrast in moving objects, such as the free-floating fibers. For the presented images, a maximal projection of the entire video was used.

Fiber Airstream Simulations

We conducted axisymmetric aerodynamic simulation with COMSOL 5.3a with v2-f turbulence model. Default model parameters for air are used. The spinneret is simulated with its physical size without rotation, the air jet in the center is simplified as a steady uniform air flow through a circular opening in the center of the spinneret. The air flow speed is set to 5m/s. The external boundaries are set to be sufficiently far away from the jet so that they do not affect the result showing in the figure.

Scanning electron microscopy (SEM)
All samples were sputter coated with 10 nm of platinum/palladium (Pt/Pd) using a Quorum Sputter Coater (EMS 300T D, Quorum Technologies) to reduce charge accumulation. Then samples were imaged with a field emitting electron microscope (FESEM SUPRA 55, Zeiss) at a voltage of 5 kV.

Fiber Diameter Measurements
SEM images (N=2, 3 FOV/sample) were imported into ImageJ (version 1.53e) for analysis. Fiber diameters were measured using ImageJ's built-in line measurement tool. The diameters of 100 representative fibers were measured in each field of view. All fiber diameters were aggregated to yield the fiber diameter distribution presented for each different fiber type.

Fiber Orientation Analysis
To visualize the fiber orientation, a square area is cropped from the SEM image and a Fast Fourier transform (FFT) was performed using NIH ImageJ. For orientation order parameter (OOP) analysis, SEM images (N=3, 3 FOV/sample) were analyzed using ImageJ's OrientationJ plugin to create color mapped images according to fiber orientation. The color mapped images were then processed through a customized MATLAB script (version 2021a, The Mathworks, Inc., Natick, MA), as previously described (*26*), that calculates each image's orientation order parameter. All calculated values fall between 0 and 1, where 0 represents a completely random, isotropic fiber sheet and 1 represents a perfectly aligned, anisotropic fiber sheet.

Fiber Production Rate
The fiber production rate was measured by depositing PCL fibers onto a 10 mm steel bar for a set duration, and then comparing the weight before and after deposition. Here a 6% w/v PCL solution was used, with a solution flow rate of 2 mL/min. The spinneret used contained three orifices, which were 400 µm each. For these experiments, the spinneret rotation rate maintained was maintained at 10,000 rpm, and the fiber stream was set to an exit pressure of 0.2 MPa. This experiment was repeated five times, with an average value of 0.1 g/min, or 0.03 g/min/orifice observed.

X-ray micro computed tomography (µCT)
X-ray micro computed tomography (µCT) imaging was completed on the Xradia 620 Versa 3D X-ray Microscope (Zeiss, Pleasanton, California, USA). The fiber sample was specially prepared on a low contrast stiff Nylon MicroSwab (TX730, ITW Texwipe) without any contrast agent staining. µCT scanning was performed using a transmission tungsten X-ray source with tube voltage of 50 kV (photon energy levels ranging from 0 to 50 keV) and current of 90 uA. 4501 projection images were captured per sample on a 16-bit 2048 × 2048 20X objective detector with achievable voxel resolutions of 0.54 µm (sample type dependent). Total scanning time for each imaging is around 20 hours with 12 seconds for each exposure.

Rat heart isolation and cryosectioning
All animal experiments for the ventricular tissue immunostaining were approved by the the Institutional Animal Care and Use Committee (IACUC) at Harvard University. Briefly, male Sprague Dawley rats (350–500 g) were anesthetized via inhalation of isoflurane (5% in $O_2$) and euthanized by cervical dislocation. The heart was removed and placed in cold (4°C), oxygenated

($95\%O_2$:$5\%CO_2$) Krebs buffer containing 10 mU/mL insulin and the following (mM): NaCl, 112.0; KCl, 4.7; $KH_2PO_4$, 1.2; $MgSO_4$, 1.2; $CaCl_2$, 2.5; $NaHCO_3$, 25.0; dextrose, 11.0 (pH 7.4). Prior to cryosectioning and immunostaining, the hearts were stored in PBS + 30% sucrose solution overnight at 4 °C, then transferred to 50% sucrose/50% optimal cutting temperature (OCT) watersoluble blend of glycols and resins for 24 h at 4 °C. They were then transferred to cryosectioning containers in 100% OCT and stored at 4 °C for 48 h. Samples were frozen by partial immersion in 2-methylbutane which was, itself, partially immersed in liquid nitrogen. Frozen hearts were stored at - 80 °C until cryosectioning by microtome (Leica). We obtained 10 µm-thick cross-sections that were transferred to glass microscope slides (Superfrost microscope slides, Sigma) and maintained at room temperature for 2 h prior to storage at -80°C until staining. For imaging, we selected cross-sections of the heart at the equator, half-way from the base to apex, where cross-sections included left and right ventrical walls but did not include heart valve features.

<u>NRVM cell and tissue culture</u>

Neonatal rat ventricular myocytes (NRVMs) were isolated from two-day-old neonatal neonatal CRL: CD (Sprague-Dawley, SD) rats. NRVMs were seeded at a density of five to seven million cells per ventricle. The NRVMs were cultured on the scaffolds with M199 culture media supplemented with 10% heat-inactivated fetal bovine serum (FBS), 10 mM HEPES, 0.1 mM MEM nonessential amino acids, 20 mM glucose, 2 mM L-glutamine, 1.5 µM vitamin B12, and 50 U/mL penicillin. Samples were incubated at 37 °C and 5% $CO_2$. At 48h post seeding the media was exchanged with M199 media containing 2% FBS and was exchanged again every 48h before testing.

The isolation of neonatal rat ventricular cardiomyocyte (NRVM) was performed using protocols was approved by Institutional Animal Care and Use Committee in Harvard University. Here, ventricles were surgically removed from 2-day old neonatal Sprague-Dawley rat pups (Charles River Laboratories, Wilmington, MA). The tissue was sectioned into ~8 equal segments and rinsed in Hanks' balanced salt solution (HBSS; Thermo Fisher, Waltham, MA), followed by digestion in 1 mg/mL trypsin (MilliporeSigma, St. Louis, MO) in HBSS solution for 13 hours under gentle agitation using a rocker. Temperatures were maintained by placing the tissue segments within an ice bath, that was placed inside a 4ºC refrigerator. Next, the tissue was further homogenized in 1 mg/mL collagenase (MilliporeSigma, St. Louis, MO) in HBSS solution at 37ºC using a series of four dissociation steps. Ventricular cardiomyocytes were then collected by centrifugation and filtered through a 40 µm cell strainer, removing any undissociated clumps. Fibroblasts were then preferentially selected against, using two 45-minute pre-plating, where fibroblasts preferentially adhere to the substrate surface, while cardiomyocytes remain non-adherent. Cardiomyocytes were then isolated and resuspended in M199 media (Life Technologies, Carlsbad, CA) supplemented with 10% (v/v) heat-inactivated fetal bovine serum (FBS; Life Technologies, Carlsbad, CA), 50 U/mL penicillin (Life Technologies, Carlsbad, CA), 3.5 g/L glucose (MilliporeSigma, St. Louis, MO), 10 mM HEPES (Life Technologies, Carlsbad, CA), 2 mM L-glutamine (Life Technologies, Carlsbad, CA), 1% (v/v) MEM non-essential amino acids (Life Technologies, Carlsbad, CA), and 2 mg/L vitamin B12. Cardiomyocytes were then counted, fiber scaffolds were seeded at an appropriate density, typically five to seven million cells per ventricle scaffold.

hIPSC-CM culture and differentiation

Human induced pluripotent stem cells (hIPSCs) were differentiated into a cardiomyocyte lineage following a 15 day differentiation protocol (Fig S15). Here, WTC-11 wildtype hiPSCs (Coriell Institute for medical research) were maintained on GelTrex (Life Technologies) coated well-plates in Essential 8 (E8) Medium (Life Technologies), containing an E8 supplement (Life technologies) and penicillin/streptomycin (Life Technologies). During the iPSC maintenance phase, media was changed daily, and cells passaged every ~3 days, first by rinsing in phosphate buffered saline (PBS) and then dissociating cells from the surface Versene (5 min, Life Technologies). Cells were then templated in E8 media containing a Y-27632 (5 μM, Life Technologies) inhibitor for one day, prior to changing to fresh E8 media. To begin stem cell differentiation into cardiomyocytes, hIPSCs were maintained until they reached ~50% confluence, at which time, differentiation was induced by introducing the WNT agonist CHIR99021 (6 μm, Stem Cell Tech) in RPMI media (1640, GlutaMAX, Life Technologies) containing a B27 supplement (minus insulin, Life Technologies) for 48 hours. Next, samples were rinsed with PBS and then changed into fresh RPMI media for 24 hours, before introducing the WNT/ β-catenin inhibitor IWR-1 Endo (5 μM, Stem Cell Tech) for another 48 hours. After this time, cells were maintained in RPMI media, changed every 48 hours. On days 9-12 of differentiation, hIPSC derived cardiomyocytes (CMs) were purified by first weaning the cells onto sodium lacatate (5 mM, Sigma-Aldrich), then introducing RPMI media containing sodium lactate (5 mM) in the absence of the B27 supplement. After 15 days of differentiation, cells were either maintained using RPMI media containing B27, which was replenished every 48 hours, or by dissociating cells using room temperature StempPro Accustase (Life Technologies). For plating, cells were maintained in STEMdiff Cardiomyocytes Support Medium (Stem Cell Tech), containing penicillin/streptomycin (Life Technologies) for the first 24 hours, before being replaced with RPMI media containing B27, which was replenished every 48 hours thereafter. Experiments were typically conducted between 7-14 days after hIPSC-CM replating.

Immunostaining

For immunostaining, 10 μm-thick cross-sections of heart or fiber scaffold with NRVM were permeabilized in 0.5% Triton-X100 for 20 min in PBS at 37 °C, followed by 2 h incubation with 1:200 dilutions of mouse anti-sarcomeric a -actinin monoclonal primary antibody (Sigma-Aldrich, clone EA-53, catalogue number A7811-11UL). Samples were then washed and concurrently incubated with 1:200 dilutions of DAPI (Sigma-Aldrich), phalloidin conjugated to Alexa-Fluor 488 (Invitrogen) and goat anti-mouse secondary antibody conjugated to tetramethylrhodamine for 2 h at room temperature. Imaging was performed using a Zeiss LSM 5 LIVE confocal microscope with a Plan-Neofluar 20× /1.3 oil objective.

Optical mapping experiments

Calcium activities of the engineered 3D fibers structure were monitored with a calcium indicator, X-Rhod-1 (Invitrogen, Carlsbad, CA), using a modified tandem-lens macroscope that (Scimedia, Costa Mesa, CA) equipped with a high-speed camera (MiCAM Ultima, Scimedia, Costa Mesa,

CA), a plan APO 0.63× objective, a collimator (Lumencor, Beaverton, OR), and a 200mW Mercury lamp (X-Cite exacte, Lumen Dynamics, Canada), and a high-spatial resolution sCMOS camera (pco.edge, PCO AG) and 880 nm darkfield LED light were incorporated into the system. For $Ca^{2+}$ imaging, an excitation filter with 580/14 nm, a dichroic mirror with 593 nm cut-off, and an emission filter with 641/75 nm (Semrock, Rochester, NY) were used. For dark field imaging, a dichroic mirror with 685 nm cut-off and long pass emission filter with 664 nm cut-off filter (Semrock, Rochester, NY) were added. Samples after 7 days culture were incubated with 2 µM X-Rhod-1 (Invitrogen) for 60 min at 37 °C, rinsed, and incubated in dye-free media for an additional 15 min at 37 °C before recording. Prior to recording for the experiments, the culture media was replaced with Tyrode's solution (1.8 mM $CaCl_2$, 5 mM glucose, 5 mM Hepes, 1 mM $MgCl_2$, 5.4 mM KCl, 135 mM NaCl, and 0.33 mM $NaH_2PO_4$ in deionized water, pH 7.4, at 37°C). The tissues were stimulated with a pulse generator. Point stimulation was applied using two platinum electrodes (Sigma-Aldrich) with 1 mm spacing with 12V amplitude and 10 ms duration. The platinum electrodes were located at 1.0 mm from the edge of "sample. For each recording, $Ca^{2+}$ and dark field images were acquired with 400 frames at a frame rate of over 10 s. Post-processing of data was conducted with custom software written in MATLAB (MathWorks) and MiCAM imaging software (MiCAM Ultima, Scimedia, Costa Mesa, CA). A spatial filter with 3 × 3 pixels was applied to improve the signal-noise ratio. Activation time of each pixel was calculated at the average maximum upstroke slope of multiple pulses of X-Rhod-1 signals over a 10 second recording window.

Particle Imaging Velocimetry Measurements

Particle Imaging Velocimetry Measurements (PIV) measurements were conducted on a Zeiss Discovery.V12 stereomicroscope, with an HBO 100 fluorescent light source, using a Basler electric ACA2500-14UC USB 3.0 camera, recorded at a 10x magnification, at a frame rate of 30-50 frames per second, with resolution of 1920x1080 pixels. Prior to recording, samples were rinsed in fresh Phosphate Buffered Saline (PBS, 37° C) and then placed in an imaging solution, comprised of Tyrode's Salt solution containing 10 µg/mL of suspended 1-5 µm green fluorescent beads (FMG – Green Fluorescent Microspheres, Cospheric, Santa Barbara, CA, USA). Samples were field stimulated with an IonOptix myopacer, using two parallel platinum electrodes positioned ~30 cm apart, at a pacing frequency of 1-2 Hz, and 10-20 volts.

Velocity map reconstruction was done in python using custom software, based on the open source package OpenPIV (*36*). Fluorescent images were prepared for analysis using a rolling ball background subtraction (NIH ImageJ). Digital image cross-correlation was then performed on a frame-by-frame basis using OpenPIV, resulting in time evolved 2D velocity vector fields. Here outliers were filtered using spatial averaging. Any vector fields which exceeded 10x the standard deviation of the local vectors (3x3 grids), were replaced using the average value of the remaining vectors. This resulted in a time dependent velocity profile, which was phase averaged over the entire cycle to smooth out local fluctuations in performance. To phase average velocity fields, the sum of the longitudinal velocities was acquired (**Fig S.12E**), and fit using a least-squares regression to a sin wave. This provided the wavelength and phase-shift of each time series. Velocity vectors were then averaged over the full-cycle, providing a single averaged cycle. The resulting averaged velocity vectors were used to perform subsequent analysis.

Cardiac Output and Ejection Fraction

As a simple metric pumping behavior, we measured the instantaneous mass flux, $N_{flux}$, passing through the basal opening of the ventricle, given by:

$$N_{flux} = \rho_f \int_{\partial S} u \, \partial l \qquad \textbf{(S2)}$$

Where $\rho_f$ is the fluid density (taken as ~978 kg/m$^3$ for Tyrode's solution), $S$ is the region of analysis surrounding the opening, $\partial S$ is the boundary of that region, $\partial l$ is a line element of $\partial S$, and $u$ is the fluid velocity (**Fig. S12B)**. Cardiac output, Co, or 2D mass transfer, was then determined by integrating the mass flux over the systolic period, and is given in units of g/m. This returns a 2D metric, which by integrating over the whole z-depth of the ventricle opening, can be used to recover the total mass transferred, given in units of g.

To estimate the ejection fraction of each ventricle, we can integrate the 2D cardiac output over the entire basal opening, convert this to a fluid volume using the fluid density and then normalize this value by the total volume of the ventricle. This model assumes even ejection velocities across the depth of the ventricle, which were approximated using measurements taken from a mid-depth. Ventricles were approximated as half-oblate ellipsoids, with their volume, V, being given by

$$V = \frac{4}{6}\pi \cdot a \cdot b \cdot c \qquad \textbf{(S3)}$$

where a,b,c are the radii of the major, and two minor axis respectively (i.e. ½ the length, width and height). Here the average length and width of ventricles after swelling were used, yielding values for a (~20 mm) and b (~6 mm) directly, with c assumed to be ½ the value of the other minor axis, given the ventricles tendency to relax. Using dimensional analysis and integration over the total cardiac output yields an estimate for the ejection fraction (EF) such that,

$$EF = \frac{\pi}{4V\rho_f} Co \cdot 2c \qquad \textbf{(S4)}$$

where the π/4 term corrects for the area or the ventricle opening, which was assumed to be a circle inscribed within a square.

Ventricle Twist

To measure rotational changes during contraction, fiber ventricle scaffolds cultured were first cultured with NRVM cardiomyocytes for 5-6 days before being at the base to a supporting rod (**Fig. S10**). Scaffolds were then suspended in a glass scintillation vial, containing ~20 mL of fresh Tyrode's solution (37 °C). Cells were stimulated using two platinum electrodes placed on either side of the ventricle, ~25 mm apart, which were connected to an IonOptix myopacer, at a pacing frequency of 1 Hz, and 10 volts. The ventricle's shape was then captured from the bottom up, through the flat bottom of the scintillation vial, using Nikon D750 camera at a pixel resolution of 6016x4016 (**Fig. S10B**). For ease of reference, green fluorescent fiducial markers were placed on the apex of the ventricle via manual pipetting (Green Fluorescent Microspheres, Cospheric, Santa Barbara, CA, USA). To estimate shifts in rotation, videos were first thresholded using NIH's ImageJ, and then the circumference of the ventricle's surface was approximated using an elliptical fit (**Fig. S10C**). This produced major and a minor axis for the ventricle, which could be used to estimate the angle or rotational orientation. Normalizing these orientations by the ventricles baring during diastole, the maximum and minimum rotation were noted throughout contraction (**Fig. S10D**), using the difference in these values to calculate the total rotational shift (**Fig. S10E**). This process was repeated for both circumferentially and helically aligned scaffolds, yielding statistically different total rotations for each alignment ($p<0.005$, $n=7$ for each condition).

Catheter Experiments

Catheter experiments were performed following previously reported protocols (*24*). Briefly, intraventricular pressure and volume (PV) were recorded simultaneously on a Millar MPVS Ultra single segment foundation system using a SPR-869NR rat PV catheter (ADInstruments). Here, ventricles were transferred to a 35 mm petri dish containing Tyrodes solution, with temperatures maintained at 37 °C using a stage top heating element (Warner Instruments). To perform PV experiments, ventricles were sutured to an ~2 mm silicone tubing, through which the catheter was inserted. Using a manufacturer-supplied acquisition systems (Millar MPVS Ultra, ADInstruments LabChart), data was acquired at a sampling rate of 2,000 Hz, and was exported for post-processing (LabChart), which was performed using a custom python script (python 3.6.5).

In post-processing, pressure and volume data from a 4.5 second interval was smoothed using a 3$^{rd}$ order univariate spline fit, and background drift was subtracted using a linear regression. To obtain a pressure-volume loops, values were phase averaged over the duration of the recording, providing a single average pressure-volume measurement.

Prior to each experiment catheter calibration was performed using a series of different sized wells drilled into an acrylic block, with known dimensions. Experiments were performed on four occasions with seven total ventricles, with one example PV loop given, however due to the low dielectric constant of gelatin, there was poor consistency across samples.

Strain Mapping, Contraction Area and Axial Shortening

To measure strain, axial deformation and changes in ventricle area during contraction an analogous 3D process to traction force microscopy (TFM) was used, with fluorescent beads serving as fiducial markers. To achieve this, ventricle scaffolds were placed on their side in a 35 mm petri dish, and were soaked in a solution of green fluorescent beads (10 µg/mL, 1-5 µm diameter, FMG – Green Fluorescent Microspheres, Cospheric, Santa Barbara, CA, USA) suspended in a Tyrode's solution (37 °C) for 20-30 minutes. This allowed for non-specific adhesion of the fluorescent beads to the scaffolds surface. Scaffolds were then rinsed, and moved to a fresh bath of Tyrode's solution containing no fluorescent beads. Samples were then field stimulated at a pacing frequency of 1-2 Hz, and 10 volt, using two parallel platinum electrodes positioned ~30 cm apart which were connected to an IonOptix myopacer system. Changes and deformations were recorded on a Basler electric ACA2500-14UC USB 3.0 camera, connected to a Zeiss Discovery.V12 stereomicroscope, with an HBO 100 fluorescent light source. Experiments were conducted at a 7.0x magnification, at a frame rate of 30-50 frames per second, with resolution of 1920x1080 pixels.

To recover strains, the open source package OpenPIV (*36*) was used, which employs digital image cross-correlation to measure displacements in corresponding images. Here we used this to measure deformations from a given reference image, in this case one frame selected during diastole for each ventricle. This generated deformation maps across the ventricle's surface, which were then normalized by the total ventricle size to produce strain values. Background pixels were filtered based on intensity to remove null frames in the deformation map. Outliers were filtered using a rolling kernel density estimate (kernel size 3x3), where the standard deviation for each kernel was measured, and local values that exceeded 5σ were replaced with mean kernel values. To recover changes in axial deformation and area, these fluorescent videos of ventricle contraction were made binary using a Huang threshold in NIH ImageJ, producing an image stark containing outlines of the ventricle's profile. Total area was measured, and the border was approximated using an elliptical fit of the ventricle's shape. This provided approximate measurements for the ventricle major and minor axis. To control for differences in contractile strength between samples, change in the major and minor axis lengths, were then normalized by the total area change that occurred for an individual ventricle, resulting in a normalized metric of basal and axial shortening (**Fig. S8D**).

Statistical Methods

Unless otherwise noted, statistical comparisons were made using a two-tailed pair wise student T test, assuming equal variance, where significance was assumed for $p$ values $< 0.05$. All error bars are given as the standard deviation unless otherwise noted. Box and whisker plots are given for the first and third interquartile range, with the mean value noted.

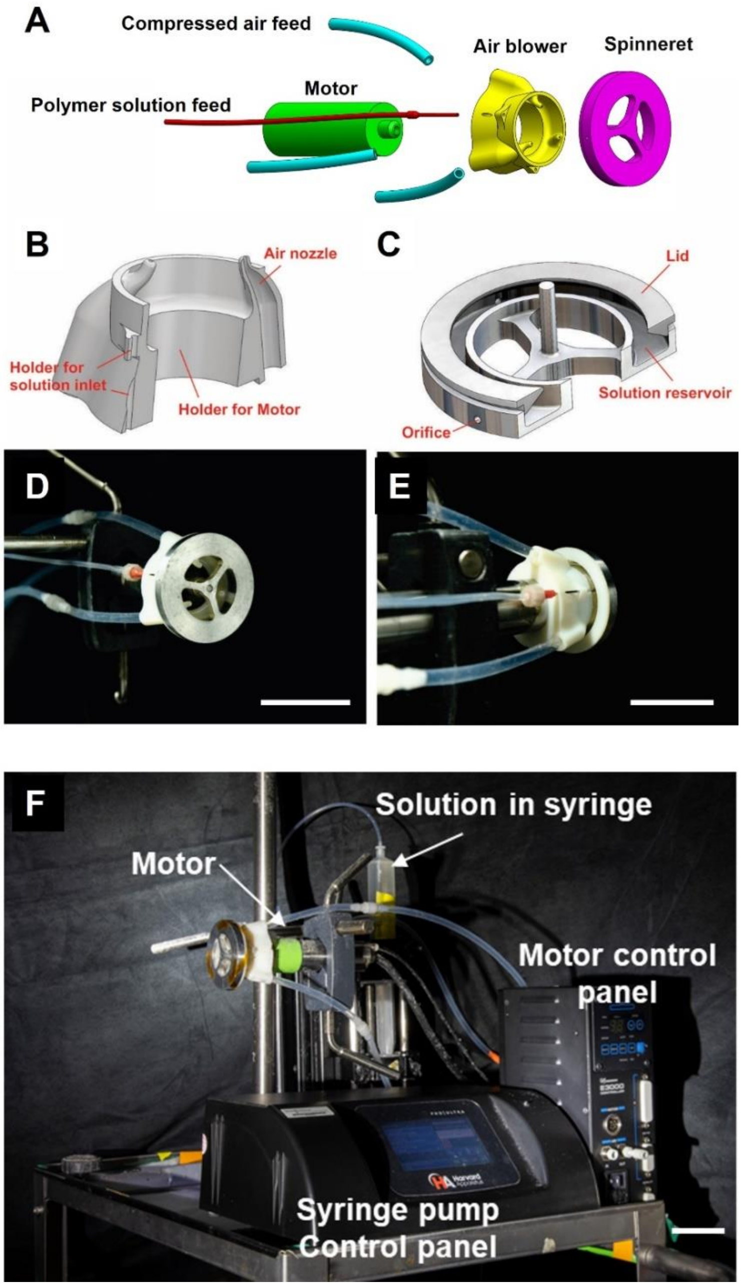


**Fig. S1 Design of the Focused Rotary Jet Spinning Platform** (**A**) Schematic of the components. The spinning platform consists of a spinneret, an air blower, a motor, and tubes that feed the polymer solution into the spinneret. Compressed air is fed into the air blower, while the spinneret is hollowed out in the center to allow air to blow out. The air blower has three nozzles, with the three streams of air merging into a single air jet in front of the spinneret. The number of air nozzles matches the number of spokes on the spinneret so that as the spinneret spins the air jet pulsates, maintaining the same directionality. (**B-C**) Cross-section of designed air blower and spinneret. (**D-F**) Photograph of the whole platform with assembled components.

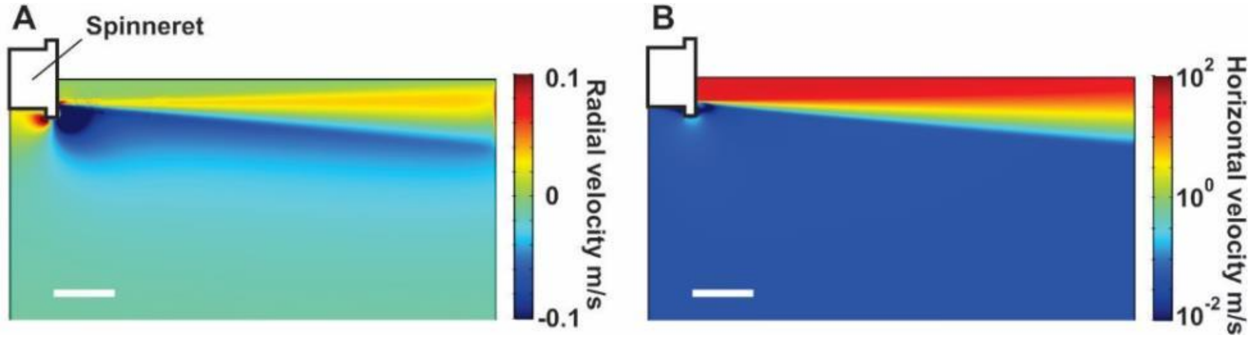


**Fig. S2 Air Jet Induced Flow Field Simulations** (**A**) COMSOL simulations showing that the jet generates a low velocity radial flow that pulls the fiber towards the air jet. (**B**) Horizontal velocity outside the air jet is orders of magnitude lower than the velocity inside the jet. At the region surrounding the spinneret where the fiber forms, the flow velocity induced by the air jet low enough such that its perturbation on fiber formation is negligible. Scale bars, 5 cm.

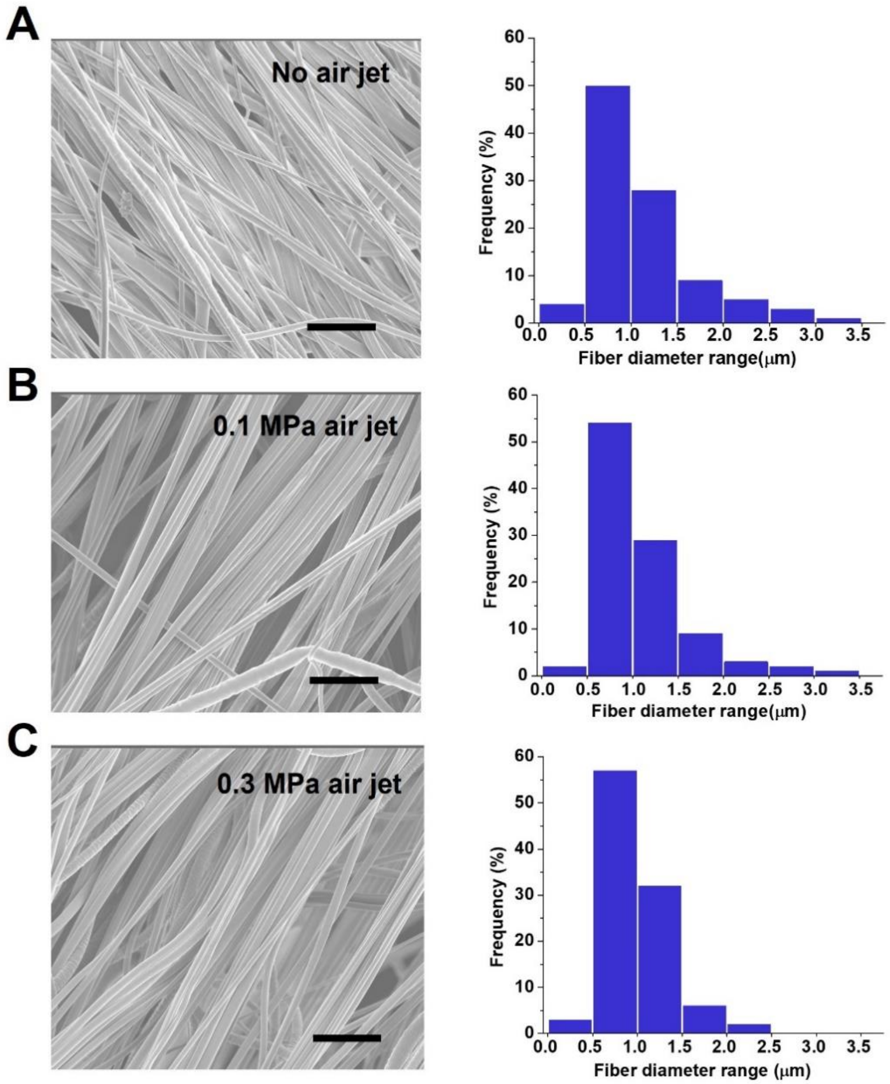


**Fig. S3 Air jet has a minor impact on fiber formation.** (**A**) Fiber collected without air jet. (**B**) Fiber collected with air jet formed under 0.1MPa compressed air. (**C**) Fiber collected with air jet formed under 0.3MPa compressed air. The fiber size distribution shifts very little under the three conditions. For each condition, 3 different samples are tested. 3 field of views are analyzed for each sample. Total fiber count for each condition >100. Scale bars, 5μm

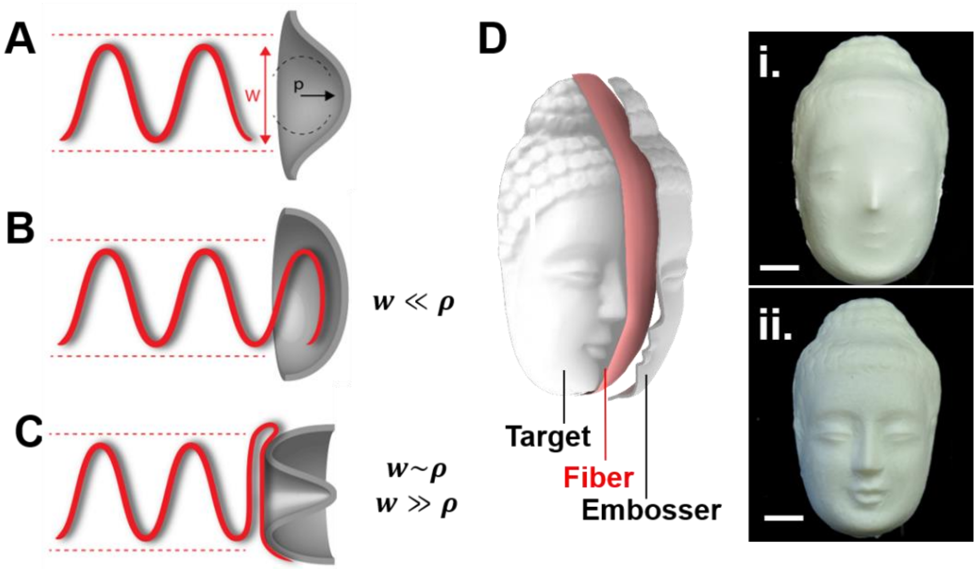


**Fig. S4 Conformity of Fiber deposition.** (**A**) The conformity of the deposition is determined by two length scales: the stream width w and the radius of curvature of the target surface ρ. (**B**) If w≪ρ the target is effectively flat for the stream, and the deposition conforms to the shape of the target. (**C**) If w~ρ or w≫ρ, overhanging fibers prevent the conformal deposition. (**D**) Deposition on a small Buddha face replica is not conformal (**i.**). Embossing forces conformity on features smaller than the stream width, resulting in a well-defined mask (**ii.**) (scale bars, 3 cm). Critical to biofabrication, conformal deposition was achieved if a feature's size was larger than the fiber stream (~5 cm) or convex in structure, while finer features could be achieved using embossing.

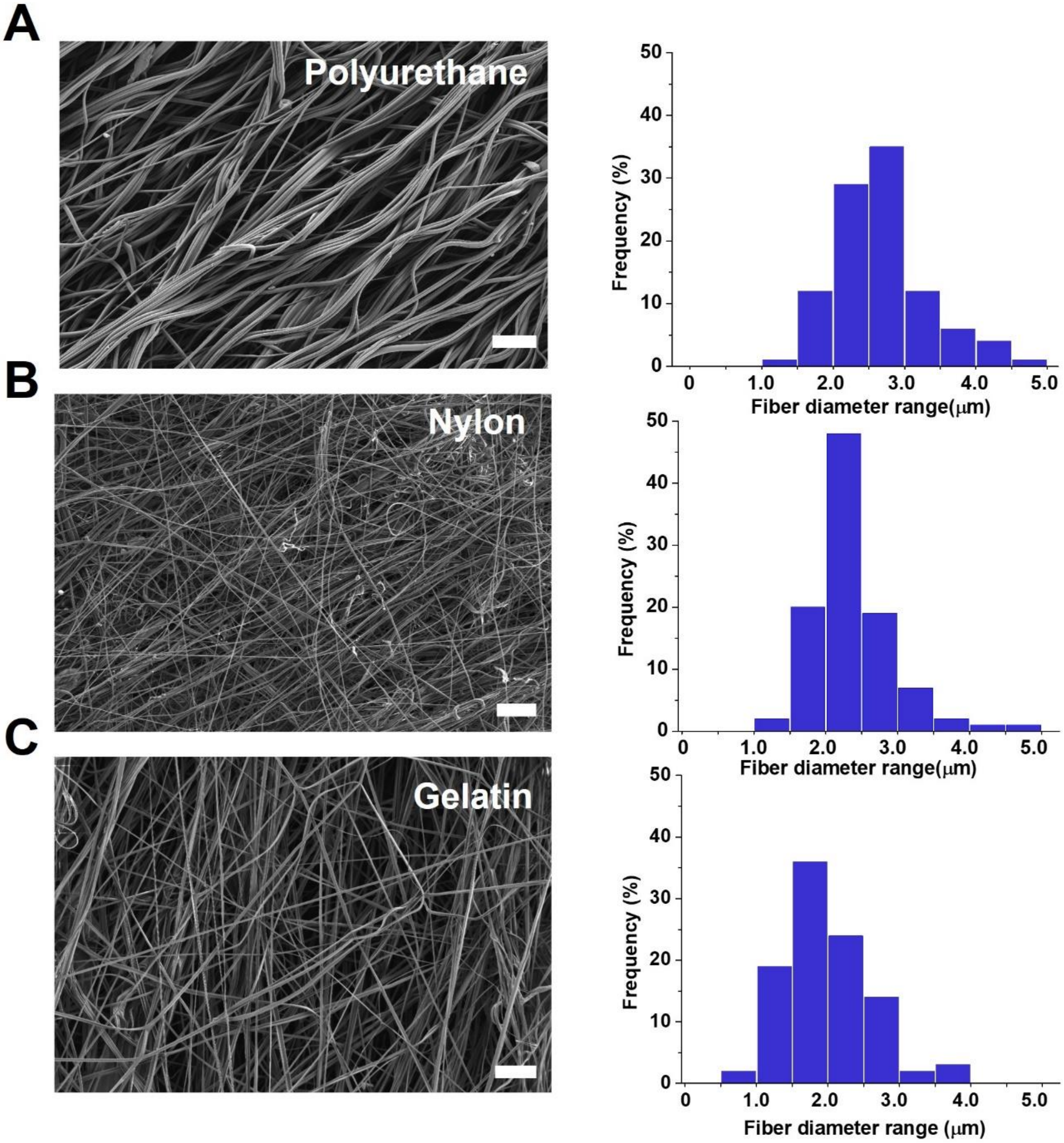


**Fig. S5 FRJS can produce a variety of polymer fibers.** (**A**) Scanning electron micrograph with corresponding histogram of fiber diameters for (**A**) polyurethane, (**B**) Nylon, and (**C**) gelatin fibers, showing that FRJS can produce a variety of fibers (Scale bars, 50 µm).

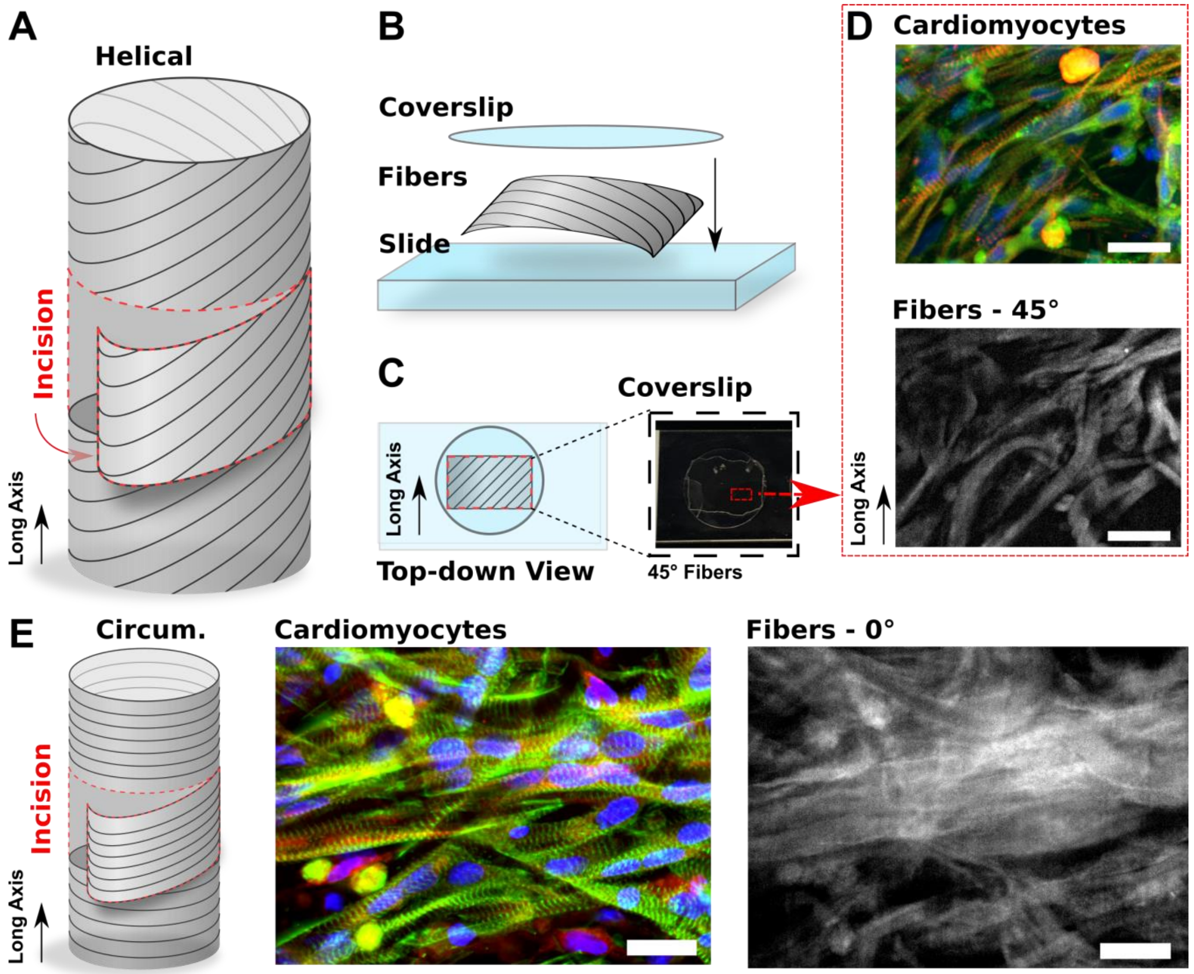


**Fig. S6 Fibers Direct Tissue Alignment and Morphogenesis. (A-b)** Schematic showing how the relative cell and fiber alignments were confirmed. Circumferentially and helically aligned fiber scaffolds cultured with NRVMs were sectioned (**A**) and mounted onto a slide (**B**), making sure to preserve the orientation of long axis. (**C**) Schematic depicting a top-down view of the mounted sample, with corresponding brightfield micrograph (α=45°). (**D&E**) Immunoflourescent staining showing cardiomyocytes are locally confluent and aligned along the direction of the fiber orientation in the tissue engineered scaffolds (upper, blue=DAPI), green=f-actin, red= α-actinin), for helically (**D**) and circumferentially (**E**) aligned fibers (Maximum Z-stack projection). Corresponding fiber autofluorescence taken from the same field-of-view (lower), showing that tissue alignment is preserved (scale bars,20 μm.). Cells were cultured for 7 days, and immunofluorescence images were background subtracted using a rolling frame average.

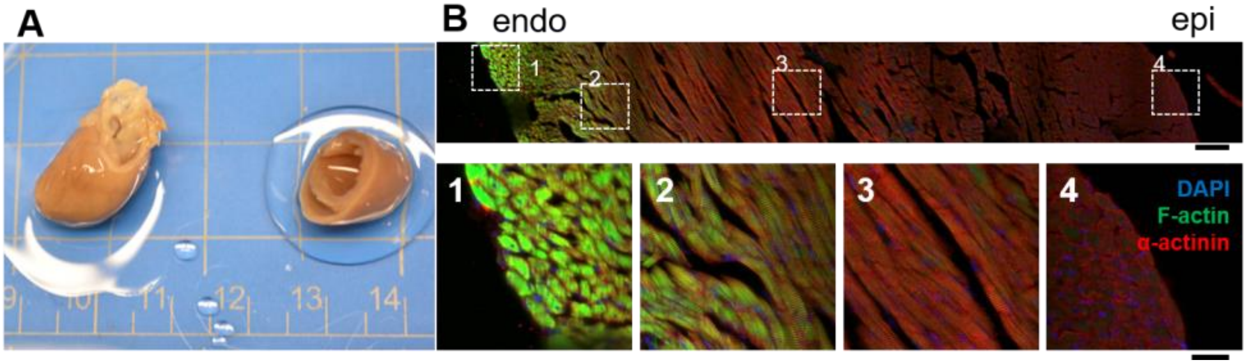


**Fig. S7 Transmural fiber orientation through an equatorial section of an adult rat left ventricle wall**. (**A**) Adult rat heart shown before (left) and after (right) it was sectioned at the equator. (**B**) Cardiomyocyte fibers in the left ventricle wall immunostained for nuclei (DAPI), F-actin (phalloidin), and α-actinin, showing fiber orientation from the endocardium (endo) to the epicardium (epi). Scale bars are 100 µm (top) and 30 µm (bottom).

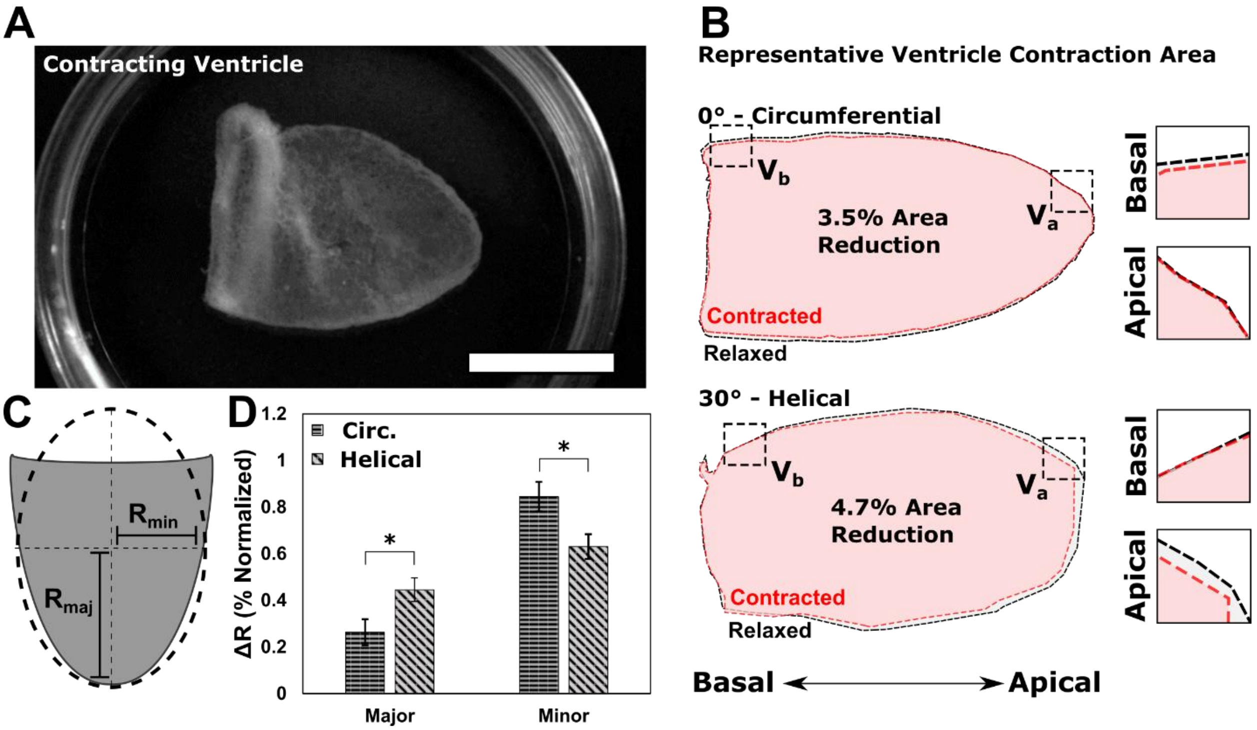


**Fig. S8 Increased Major Axis Shortening in Helically Aligned Ventricles. (A)** Photograph of a representative ventricle scaffold, visually isolated for thresholding (scale bar, 10 mm) (**B**) Representative perimeter outlines for circumferentially and helically aligned ventricles during contraction (red) and relaxation (black). Magnified insets, taken from the highlighted basal ($V_B$) and apical ($V_A$) regions, showing minimal apical and increased basal changes in the circumferentially aligned ventricles. (**C**) Schematic diagram showing quantification method of axial shorting using elliptical fitting of thresholded ventricle areas, to obtain the semi-major ($R_{maj}$) and semi-minor axis ($R_{min}$) lengths (**D**) Average change (ΔR) in in the semi-major and semi minor axis for circumferentially and helically aligned ventricles, normalized by the total area change, showing increased apical shortening in the helically aligned case. $n \geq 5$ ventricles for each test condition**.** * indicates $p<0.05$ as determined by a pairwise students T-test.

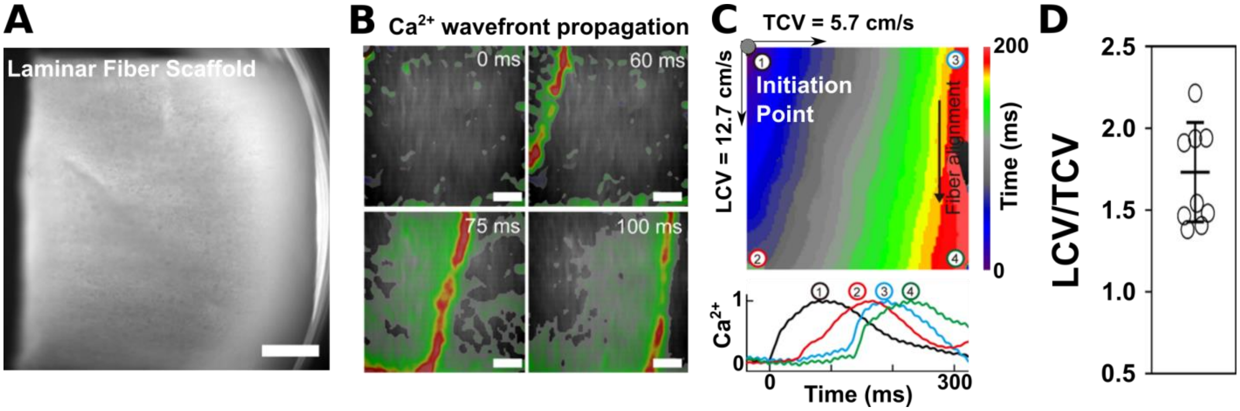


**Fig. S9 Calcium Wave Propagation in Laminar Fiber Tissues. (a)** Micrograph of laminar fiber scaffold seeded with NRVMs (scale 2 mm) (**B**) Bright field micrograph super imposed with calcium signal, showing wave front propagation at different time points after stimulation (scale 2 mm). (**C**) Isochrone of calcium wave propagation, showing the conductance velocity in the longitudinal (LCV) and transverse (TCV) direction of fiber alignments (upper). Calcium signal intensity given as a function of time for each of the four corners (lower). (**D**) Ratio of Longitudinal (LCV) to transverse (TCV) conduction velocities (n = 9 samples).

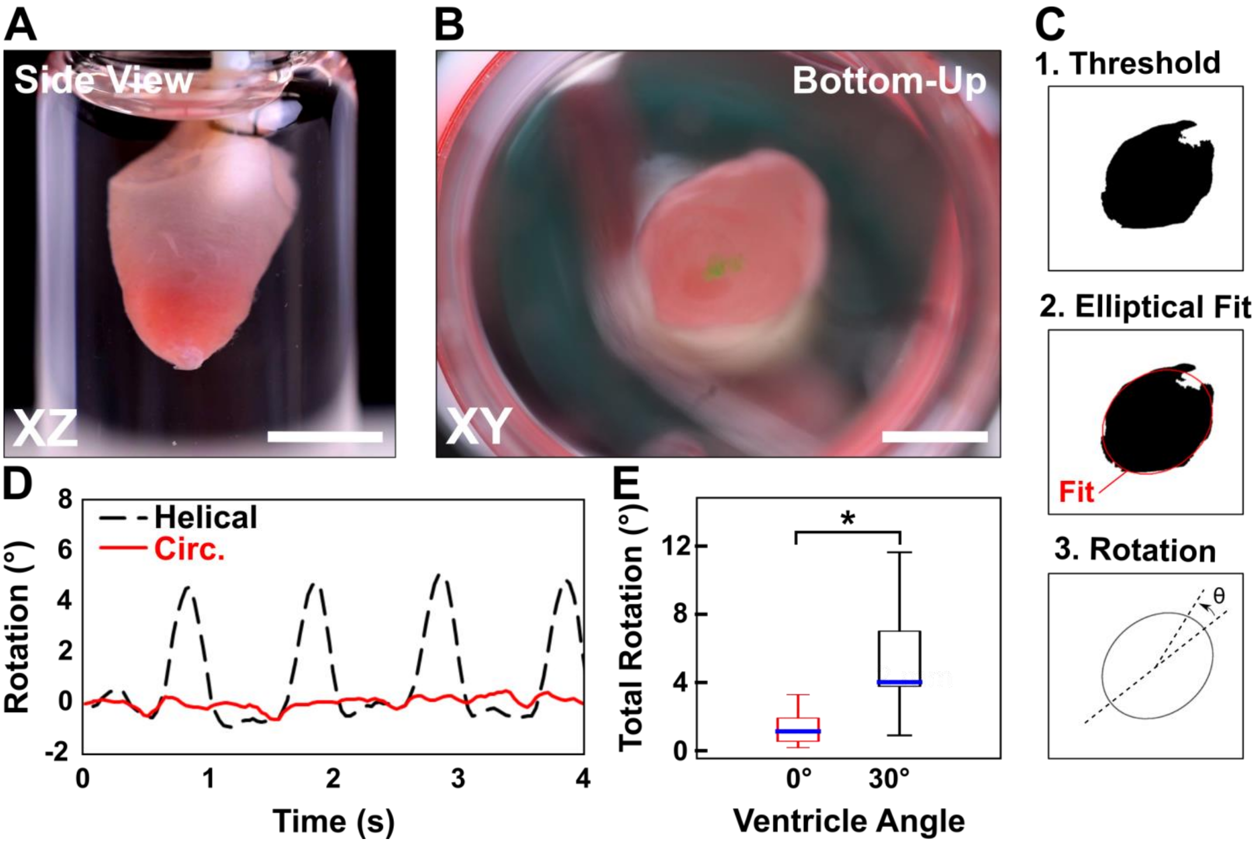


**Fig. S10 Modeling Ventricle Twist** (**A**) Ventricle scaffold sutured at the base and suspended, allowing free rotation about the apex (XZ Side view, scale bar, 10 mm) (**B**) Bottom-up view of a suspended ventricle scaffold for quantifying rotation (XY Bottom-Up View, scale bar, 5 mm) (**C**) Schema for mapping rotational displacement. Scaffolds are thresholded, and the boundary is fit to an ellipse. Changes in the angle of the ellipses major axis are recorded as rotational displacements. (**D**). Time course of rotational displacement for representative HA (α=30˚) and CA (α=0˚) ventricles, demonstrating apical twist in the HA case (1 Hz, 10v Field Stimulation). (**E**) Average rotational displacement as a function of ventricle alignment. (n = 7 ventricles each, Statistically significant difference, $p<0.005$).

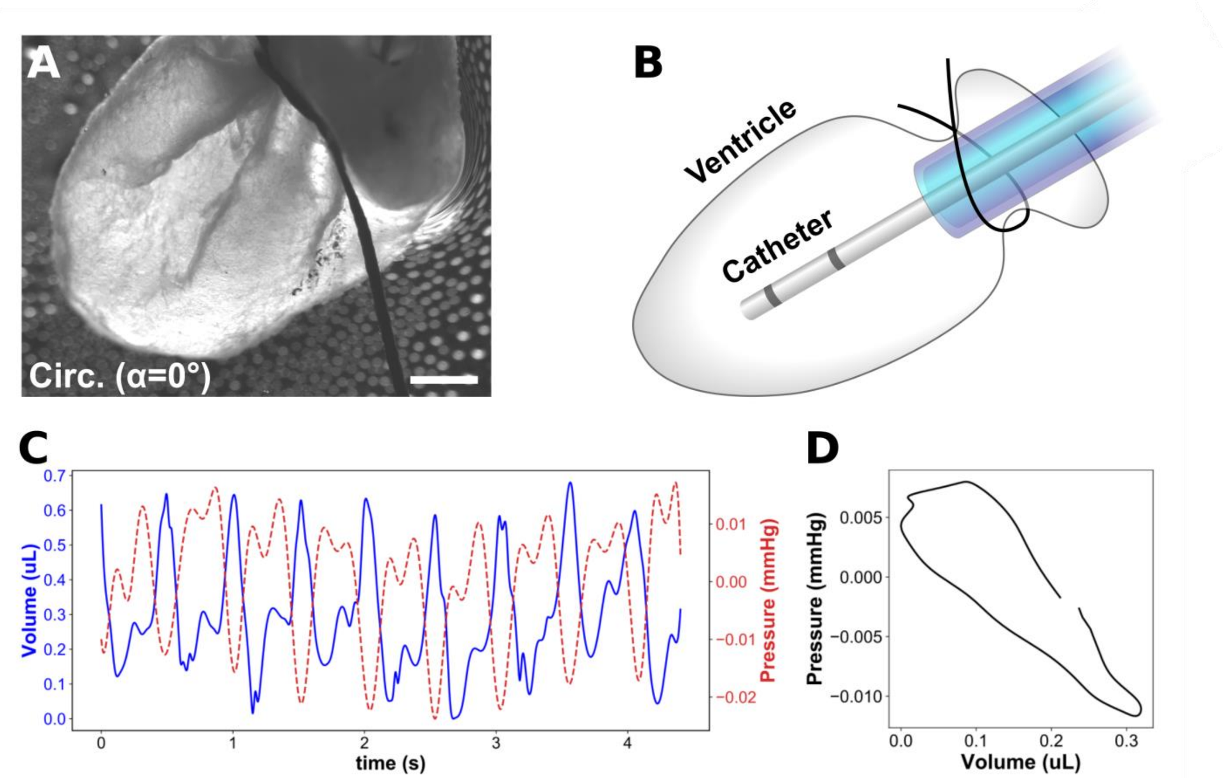


**Fig. S11 Ventricle Pressure Volume Measurements.** (**A.**) Darkfield micrograph a circumferentially aligned gelatin fiber ventricle scaffold seeded with NRVMs, sutured to a Pressure-Volume catheter. (Scale bar 2 mm). (**B**) Illustration of catheter's localization within the ventricle scaffold. (**C**) Trace of Pressure-Volume changes taken during spontaneous cardiac contraction. Values normalized to ambient conditions. (**D**) Pressure-Volume loop, time averaged over the complete trace. As the model systems did not contain valves, the resulting measurement lacked any indicators of isovolumetric relaxation, as is consistent with valvular defects.

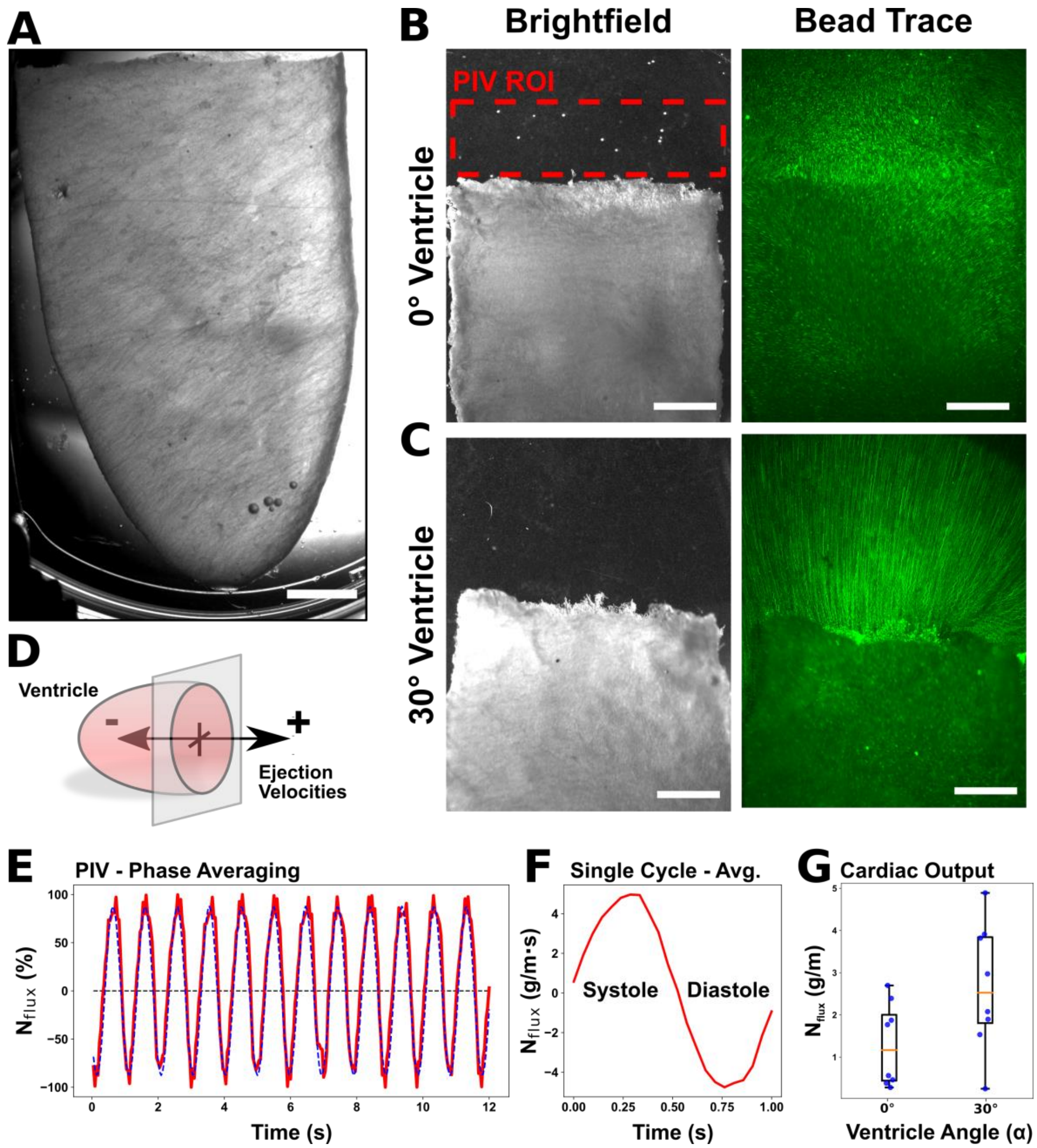


**Fig. S12 Particle Imaging Velocimetry of Helically Aligned Ventricles. (A)** Bright field micrograph of a helically aligned ventricle, resting against the base of a petri dish to prevent travel (scale bar 2 mm). Increased magnification micrograph of the basal region of circumferentially (α=0˚)(**B**) and helically (α=30˚) (**C**) aligned ventricle scaffolds (**left**), with the corresponding trace showing the flow path of fluorescent bead's (right). (Max Projection Taken Over 4 s, scale bars-2mm). Flow paths were generally increased for helically aligned scaffolds. PIV was captured over the basal region of interest (ROI) highlighted in yellow. (**D**) Schematic showing the relative direction of the fluid velocity fields. (**E**) Normalized instantaneous mass flux occurring over the highlighted ROI (red line), for a circumferentially aligned ventricle (α=0°). Mass flux was fit using a sin function (blue) to determine the period and phase of contraction. (**F**) Phase averaged instantaneous mass flux occurring over a single contraction cycle, obtained by averaging each frame over the entire collection period. (**G**) Total mass flux, or cardiac output, for circumferentially and helically aligned ventricles, obtained by integrating over the instantaneous mass flux for the systolic period.

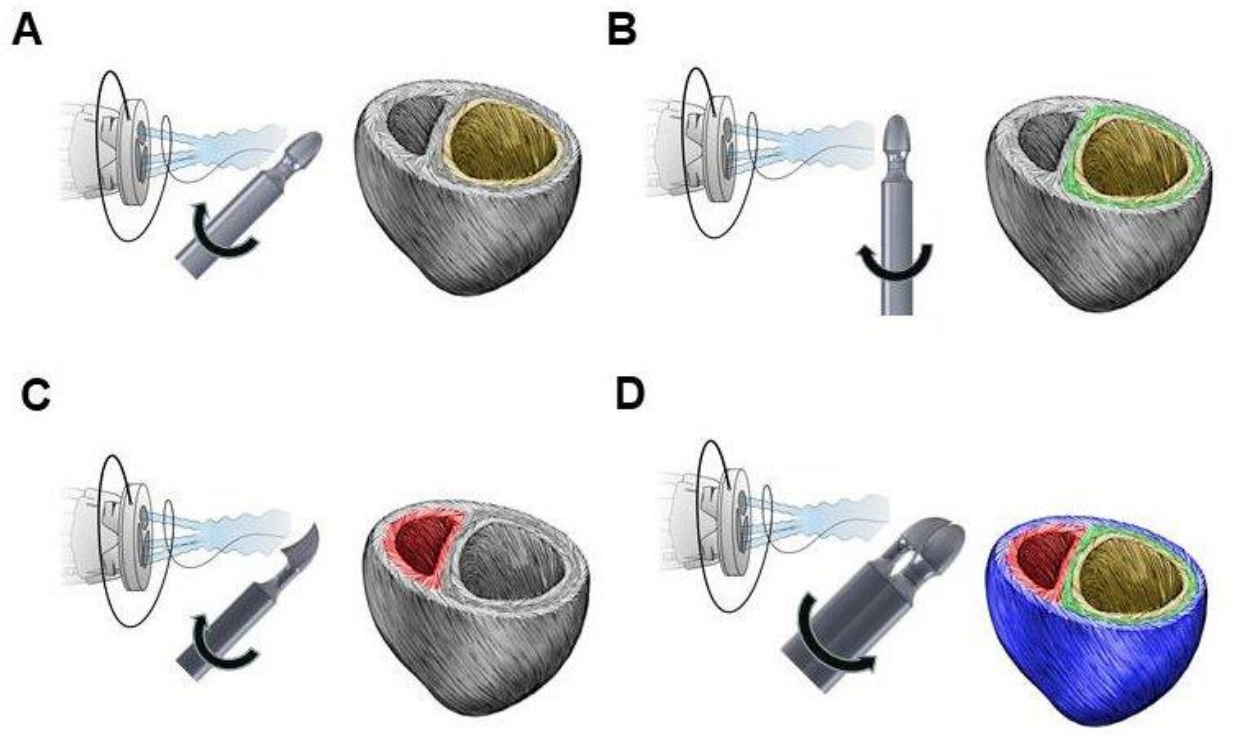


**Fig. S13 Four-step manufacturing of the tri-layered dual chambered ventricle model**. (**A**) A 3D printed target with the shape of the left ventricle is pointed into the fiber stream at 45º. The target rotates counterclockwise at 2,000RPM to collect the helically aligned inner layer. (**B**) The target is turned vertical and rotates at 10,000RPM to collect the circumferentially aligned middle layer. (**C**) A 3D printed target with the shape of the right ventricle is pointed into the fiber stream at 45º. The target rotates counterclockwise at 2,000RPM to collect the helically aligned inner layer. (**D**) The two targets with the deposited fiber are taped together and pointed into the fiber stream at 45º. The combined target rotates clockwise at 2,000RPM to collect the helically aligned outer layer.

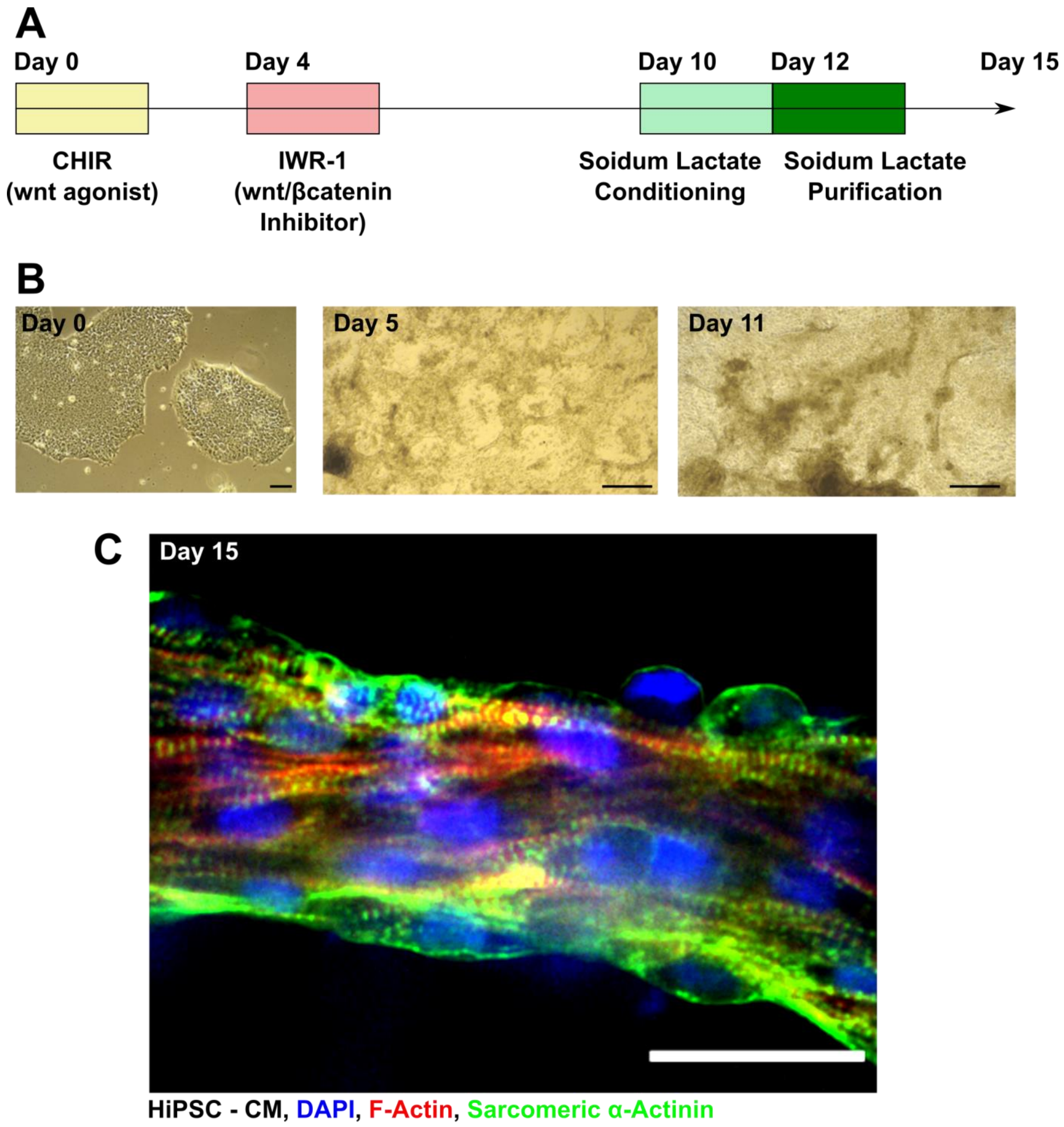


**Fig. S14 Human stem cell derived cardiomyocyte differentiation**. (**A**) Schematic overview of the fifteen day differentiation protocol, including sodium lactate purification steps. (**B**) Brightfield micrographs of stem cell differentiation, showing changes in cellular morphology at Day 0, Day 5 and Day 11 in culture. (**C**) Representative immunoflourescent micrograph of stem-cell derived cardiomyocytes after fifteen days in culture, showing the formation of aligned cardiomyocyte bundles expressing well-organized sarcomeric α-actinin (DAPI-Blue, F-Actin-Red, sarcomeric α-actinin – Green, scale bar, 50 µm).

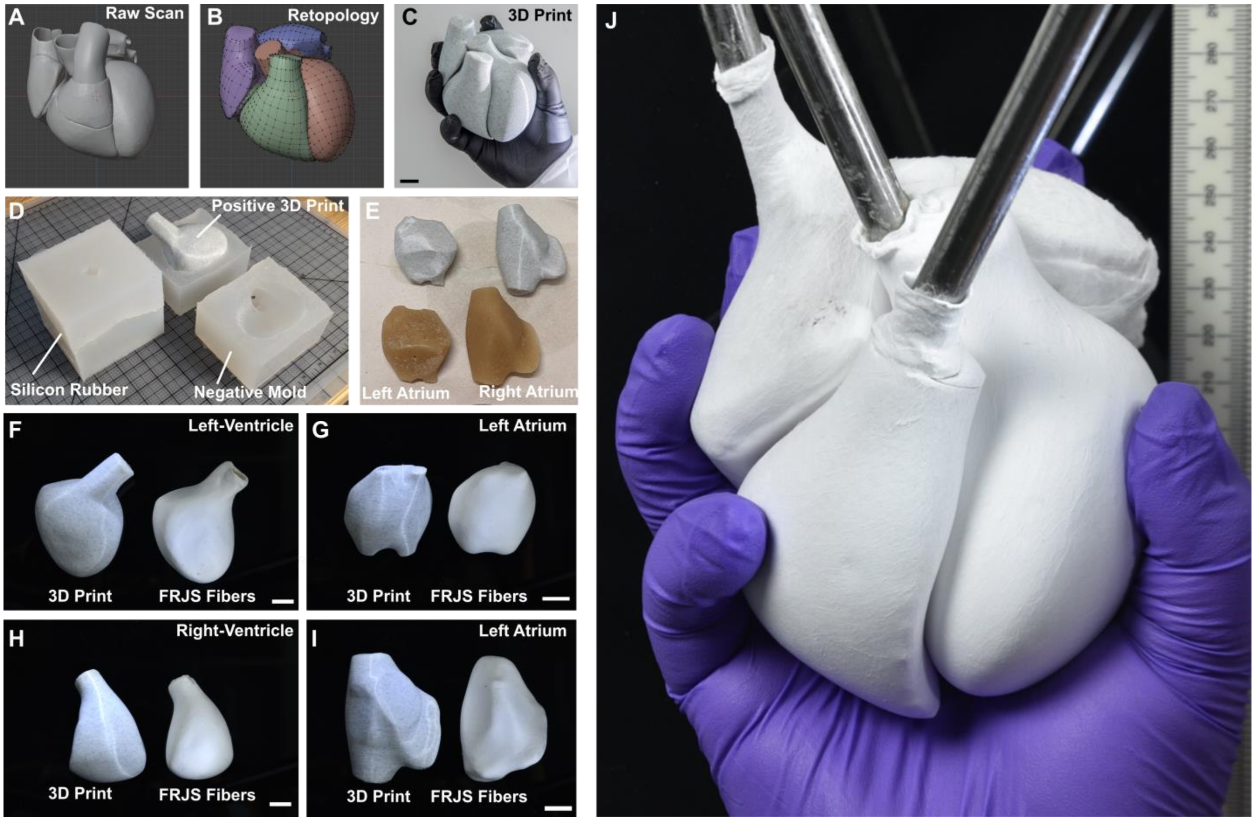


**Fig. S15 Full scale four-chambered human heart model**. (**A**) 3D model of a raw scan of the human heart. (**B**) Retopology of the human heart scan to improve printability, separated into four distinct chambers. (**C**) 3D printed model of the human heart (PLA). (**D**) Image of the 3D silicone rubber negative molds used to reproduce the heart's morphology in sugar. (**E**) Resulting sugar molds (lower) alongside 3D printed models of the heart's chambers (upper). (**F-I**) Images comparing 3D printed heart chambers (left), alongside standalone FRJS spun fiber models (PCL, right), for the left-ventricle (**F**), Left-Atrium (**G**), Right Ventricle (**H**), and Left Atrium (**I**). (**J**) Fully assembled four chamber fiber model of the human heart, still supported on sugar molds (all scale bars, 2 cm).

**Movie Captions**

**Movie S1 Fiber Stream Profile.** (**A**) Spinneret on black background producing fibers, with corresponding (**B**) Differential contrast video, showing the steam of fibers being produced. Differential contrast used to increase contrast of barely perceptible fibers.

**Movie S2 Targeted Fiber Deposition**. Video depicting the fiber deposition process onto a 3D model of the Buddha's face. After spinning, a negative mold is pressed onto the fibers, embossing the face's features. Timestamp is given, showing the total process takes under 6 minutes.

**Movie S3 Multi-layer Fiber Alignment**. X-ray micro computed tomography segmentation of a multilayered fiber block. Sectioned slices show the different fiber alignments, while moving through the block segment.

**Movie S4 Helical Fiber Alignments. (A-B)** X-ray micro computed tomography of a helically aligned fiber scaffold showing side-on (**A**) and top-down (**B**) views.

**Movie S5 Ventricle Scaffolds. (A-B)** Top-down view of helically (**A**) and circumferentially (**B**) aligned ventricle scaffolds seeded with NRVMs, showing spontaneous contractions five days after initial culture. (**C**) Side-on view of a helically aligned ventricle scaffold being field stimulated, showing paced contractions (1 Hz, 20 V).

**Movie S6 Ventricle Strain Mapping. (A-D)** Real time fluorescent micrograph of circumferentially (**A**) and helically (**C**) aligned ventricles coated in fluorescent beads undergoing field stimulation (1 Hz, 20 v), with corresponding strain maps (**B&D**) recovered using digital image cross-correlation.

**Movie S7 Alignment Dependent Calcium Wave Propagation. (A&B)** Fluorescent micrograph of calcium transience projected onto a brightfield image of a helically (**A**) and circumferentially (**B**) aligned model ventricle scaffold, with extended basal regions.

**Movie S8 Ventricle Twist. (A&B)** Bottom-up view of ventricle scaffolds suspended from a rod by their base, for helically (**A**) and circumferentially (**B**) aligned fiber scaffolds (green fiducial markers placed for visual reference). (**C&D**) Corresponding thresholded image (blue), showing how edge features are used with an elliptical fit (red), to observe rotational displacement from a starting position (white), for helically (**C**) and circumferentially (**D**) aligned scaffolds. (**E**) Graph showing the real-time ventricle twist, or rotation, for each of the given scaffolds.

**Movie S9 Particle Imaging Velocimetry Measurements.** (**A**) Brightfield micrograph of a circumferentially aligned ventricle. (**B**) Fluorescent micrograph centered on the basal region, with neutrally buoyant fluorescent beads present. (**C**) High magnification inset taken from the corresponding region of interest (ROI), after applying background subtraction. (**D**) Instantaneous mass flux, or cardiac output, occurring within the highlighted region for the corresponding circumferentially aligned ventricle and a representative helically aligned ventricle. (**E**) Phase averaged velocity map from the highlighted basal region, used to recover mass flux.

**Movie S10 Dual Chamber Ventricle (DCV) Manufacture.** Video depicting the manufacturing process for a DCV, where first the left ventricle is coated in fibers at multiple angles, and then the right ventricle is coated and adhered to the left ventricle using an additional fiber coating layer.

**Movie S11 Dual Chamber Ventricle (DCV) μCT.** X-ray micro computed tomography of the DCV's fiber structure.

**Movie S12 Dual Chamber Ventricle (DCV) Septal Wall.** High magnification X-ray micro computed tomography slice through of the fiber based DCV's septal wall.
